# Novel tensor decomposition-based approach for cell type deconvolution in Visium datasets when reference scRNA-seqs include multiple minor cell types

**DOI:** 10.1101/2025.04.18.649484

**Authors:** Y-H. Taguchi, Turki Turki

**Affiliations:** Department of Physics, Chuo University, Tokyo 112-8551, Japan; Department of Computer Science, King Abdulaziz University, Jeddah 21589, Saudi Arabia

**Keywords:** Visium, Spatial transcriptomics, tensor decomposition, deconvolution

## Abstract

We have applied tensor decomposition (TD)-based unsupervised feature extraction (FE) to integrate multiple Visium datasets, as a platform for spatial gene expression profiling (spatial transcriptomics). As a result, TD-based unsupervised FE successfully obtains singular value vectors consistent with the spatial distribution, that is, singular value vectors with similar values are assigned to neighboring spots. Furthermore, TD-based unsupervised FE successfully infers the cell-type fractions within individual Visium spots (i.e., successful deconvolution) by referencing single-cell RNA-seq experiments that include multiple minor cell types, for which other conventional methods—RCTD, SPOTlight, SpaCET, and cell2location—fail. Therefore, TD-based unsupervised FE can be used to perform deconvolution even when other conventional methods fail because it includes multiple minor cell types in the reference profiles, although it cannot be used in typical cases. TD-based unsupervised FE is thus expected to be applied to a wide range of deconvolution applications.

## 1. Introduction

Spatial transcriptomics, involving spatial measurement of gene expression is currently a popular topic [1]. In addition to the development of measurement technology, analytical technology has also been co-developed. Spatial transcriptomics, particularly the 10x Genomics Visium platform [2], has fundamentally transformed our ability to study tissue architecture by capturing both transcriptomic information and its spatial context. However, each Visium “spot” often encapsulates gene expression from multiple cells [2], complicating the direct interpretation of cell-type composition [3]. A host of computational deconvolution methods has emerged to address this issue, aiming to disentangle mixed spot-level signals into individual cellular components using single-cell RNA sequencing (scRNA-seq) references or other probabilistic and machine learning paradigms [3].

A key example is Robust Cell Type Decomposition (RCTD), which employs a probabilistic framework to robustly assign cell-type identities to spots, modeling both biological and technical variability [4]. SPOTlight adopts a seeded non-negative matrix factorization (NMF) regression strategy using scRNA-seq data to deconvolute spot-level gene expression and provide interpretable spatial maps of cell-type distributions [5]. SpaCET combines deconvolution with spatial autocorrelation analysis, illuminating local cell neighborhoods and potential interactions within tissue sections [6]. Meanwhile, cell2location uses a Bayesian approach to jointly model cell abundance and gene expression, inferring the spatial organization of fine-grained cell types across Visium spots [7].

Beyond these four applications, several methods have become integral to the spatial deconvolution toolkit. Stereoscope employs variational inference to probabilistically reconcile scRNA-seq references with spatial transcriptomics data, thus estimating the presence and abundance of different cell types at each spot [8]. Tangram leverages deep learning to align high-dimensional single-cell RNA-seq profiles with Visium spots, inferring how various cell types and subtypes map onto histological structures [9]. Additionally, integrative pipelines within widely used bioinformatics frameworks (e.g., Seurat [10–12], Giotto [13]) facilitate spot deconvolution in tandem with downstream analyses, such as trajectory inference and cell-cell interaction studies. Together, these methods provide a dynamic and continuously expanding arsenal for uncovering cellular heterogeneity and spatial organization, enabling unprecedented insight into tissue biology across both healthy and disease contexts.

On this occasion, we noticed that some of these tools fail to deconvolute Visium when the reference single-cell gene expression includes multiple minor cell types (see the later part of this study). Two kinds of single-cell gene expression are used as a reference. One is the reference for general use, that is, not specific to the targeted Visium data and another is specific to the targeted Visium data. The latter is particularly important, for example, in precision medicine intended for personal therapy. Thus, although deconvolution is expected to be accurate, especially when the reference single-cell gene expression is associated with the targeted Visium dataset, this is not the case when reference single-cell gene expression includes multiple minor cell types. To address these problems, we have developed a strategy where TD based unsupervised FE [14] has been applied to the Visium dataset. In this study, after demonstrating that four of conventional methods, RCTD, SPOTlight, SpaCET, and cell2location fail when reference single-cell gene expression includes multiple minor cell types, we have demonstrated the success of our proposed strategy.

## 2. Materials and Methods

### 2.1. Visium

The Visium dataset [15] analyzed in this study was retrieved from GEO with GEO ID GSE270382, and is composed of four Visium datasets, two of which are Tau-dKI, uninjected and two other of which are Tau-dKI, injected. They were obtained from the brains of mice in which AppNL-G-F/MAPT was doubly knocked. The downloaded matrix files were loaded into R by the `read.csv` command and formatted in a sparse matrix format using the `spMatrix` function in the Matrix package.

### 2.2. Single-cell RNA-seq and annotated cell fraction reference

Single-cell RNA-seqs [15] associated with the aforementioned Visim datasets, that is, taken from the same mouse brains, were retrieved from GEO with GEO ID GSE270385 and GSE270386. Two of the four datasets were obtained from GSE270385 and the remaining two were obtained from GSE270386 with GEO IDs GSM8341772 and GSM8341774, respectively. The .csv files were loaded into R using the `read.csv` command. We also inferred the cell types of individual cells in single-cell RNA-seq using the `SingleR` function in the SingleR package [16] in R. Annotation was performed using the MouseRNAseqData() dataset in celldex package [17]. The generated cell-type fractions are listed in Table 1.

**Table 1.** Fraction of cell-type annotation in reference single-cell gene expression associated with four Visium datasets (1 ≤ *k* ≤ 4).

| No. | $k$ | 1 | 2 | 3 | 4 |
| --- | --- | --- | --- | --- | --- |
| 1 | Astrocytes | 15 | 8 | 50 | 21 |
| 2 | B cells | 4 | 2 | 0 | 1 |
| 3 | Epithelial cells | 1 | 0 | 1 | 0 |
| 4 | Fibroblasts | 8 | 4 | 7 | 1 |
| 5 | Macrophages | 3 | 4 | 1 | 0 |
| 6 | Microglia | 3273 | 2950 | 1414 | 1697 |
| 7 | Neurons | 4638 | 3956 | 3987 | 4590 |
| 8 | NK cells | 2 | 1 | 1 | 0 |
| 9 | Oligodendrocytes | 6476 | 5365 | 3829 | 4274 |
| 10 | T cells | 30 | 16 | 39 | 11 |
| 11 | Granulocytes | 0 | 4 | 17 | 0 |
| 12 | Monocytes | 0 | 1 | 3 | 0 |
| 13 | Dendritic cells | 0 | 0 | 1 | 0 |
| 14 | Endothelial cells | 0 | 0 | 1 | 0 |

### 2.3. Detailed setups and parameters of conventional methods

For all cases, individual Visium datasets were analyzed separately in contrast to TD-based unsupervised FE, which integrates four Visium datasets (see below).

#### 2.3.1. RCTD

Function `create.RCTD` was performed by max_cores = 2, CELL_MIN_INSTANCE=1 to ensure that RCTD does not ignore the minor cell types. Then the function `create.RUN` was performed in the doublet_mode = ‘multi’ to let RCTD to attribute more than two cell types to individual spots.

#### 2.3.2. SPOTlight

SPOTlight was performed with the default settings (for more details, see the source code provided in the Github link)

#### 2.3.3. SpaCET

SpaCET was also performed with the default settings (for more details, see the source code provided in the Github link)

#### 2.3.4. cell2location

Model learning was performed using the `train` function with max_epochs=300, batch_size=2500, train_size=1, lr=0.002, accelerator=“cpu”. `Cell2location` to initialize Visium data cell type annotation was performed with N_cells_per_location=30, detection_alpha=200. Then the `train` function was performed with max_epochs=30000, batch_size=adata_vis.n_obs, train_size=1 (for more details, see the source code provided in the Github link).

### 2.4. TD based unsupervised FE

Suppose that we have tensor *x*_*ijk*_ ∈ ℝ^31053×4992×4^ which represents the *i*th gene expression at the *j*th spot of the *k*th Visium dataset. Higher-order singular value decomposition [14] (HOSVD) is applied to *x*_*ijk*_ to obtain

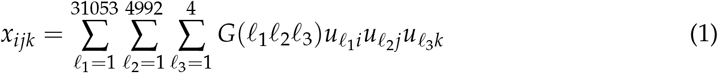

where *G* ∈ ℝ^31053×4992×4^ is a core tensor that represents the contribution of 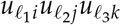 toward 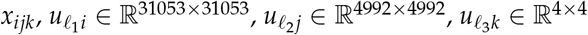 are singular value matrices and orthogonal matrices.

In this study, HOSVD was performed by applying the `irlba` function in the irlba package to unfolded matrix *x*_*ij′*_ where *j′* = *j* + 4992(*k* − 1) as

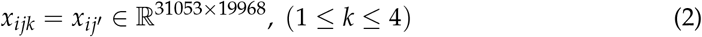

and we got

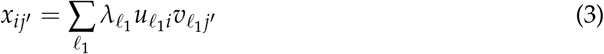

The reason why HOSVD can be replaced with SVD applied to the unfolded matrix is because HOSVD is nothing but SVD applied to the unfolded matrix.

Spots specifically contributing to the *ℓ*th singular value vector, *u*_*ℓi*_, can be selected by attributing *P*-values to the *i*th gene by assuming that the null hypothesis *u*_*ℓi*_ obeys Gaussian distribution,

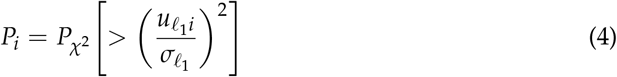

where 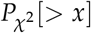 is the cumulative *χ*^2^ distribution where the argument is larger than *x* and 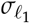 is the standard deviation. The obtained *P*-values are corrected by the BH criterion and the *i*th genes associated with adjusted *P*-values less than 0.01 are selected.

In the above computation, 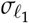 is optimized such that 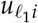 obeys Gaussian distribution as much as possible. The procedure is as follows. First, we need to compute *P*_*i*_ assuming some 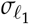. Then *P*_*i*_s are corrected with the BH criterion, and *i*th genes associated with adjusted *P*_*i*_ less than 0.01 are excluded. Next, we compute the histogram of *P*_*i*_ for the non-excluded genes and the standard deviation of the histogram, *σ*_*h*_, as

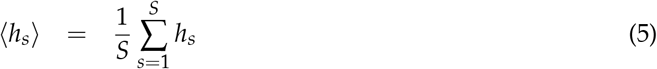

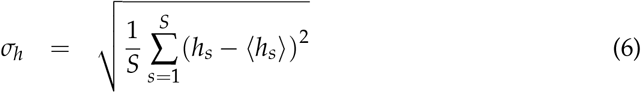

where *h*_*s*_ is the histogram of *P*_*i*_ at the *s*th bin and *S* is the total number of bins. We tune 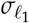 so as to minimize *σ*_*h*_.

The reason for optimizing 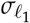 is as follows. Because 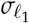 should be the standard deviation of 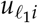, supposedly obeying the null hypothesis, Gaussian, 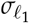 should be computed using only non-excluded genes. If we do not exclude 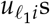 associated with smaller adjusted *P*-values, because 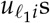 associated with smaller adjusted *P*-values have larger absolute 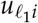 is over estimated and *P*_*i*_s are also overestimated. This results in a fewer number of selected genes. To avoid this situation, we need to compute 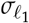 using only non-excluded 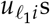.

### 2.5. Detailed procedures

To see how we can reach the results in Figs. 20, 21, and 22, we explain how we apply TD based unsupervised FE to the Visium datasets, step by step (Fig. 1). First, we apply TD-based unsupervised FE, obtain 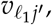 and plot *v*_2*j′*_ on the spatial coordinate (Fig. 2). Here the *j′*th spots in 1 ≤ *j′* − 4992(*k* − 1) ≤ 4992 belong to the *k*th Visium datasets. As nearby spots share the similar amount of *v*_2*j*_, i.e., consistent with the spatial coordinate, we conclude that *ℓ*_2_ = 2 is the informative and variable component. Then using *u*_2*i*_ corresponding to *v*_2*j*_, we attributed *P*-values to individual genes (*i*s) and selected 277 genes to identify genes whose expression was expected to be consistent with the spatial coordinate. We can expect these 277 genes are consistent with the spatial coordinate as well as biologically important.

**Figure 1.**
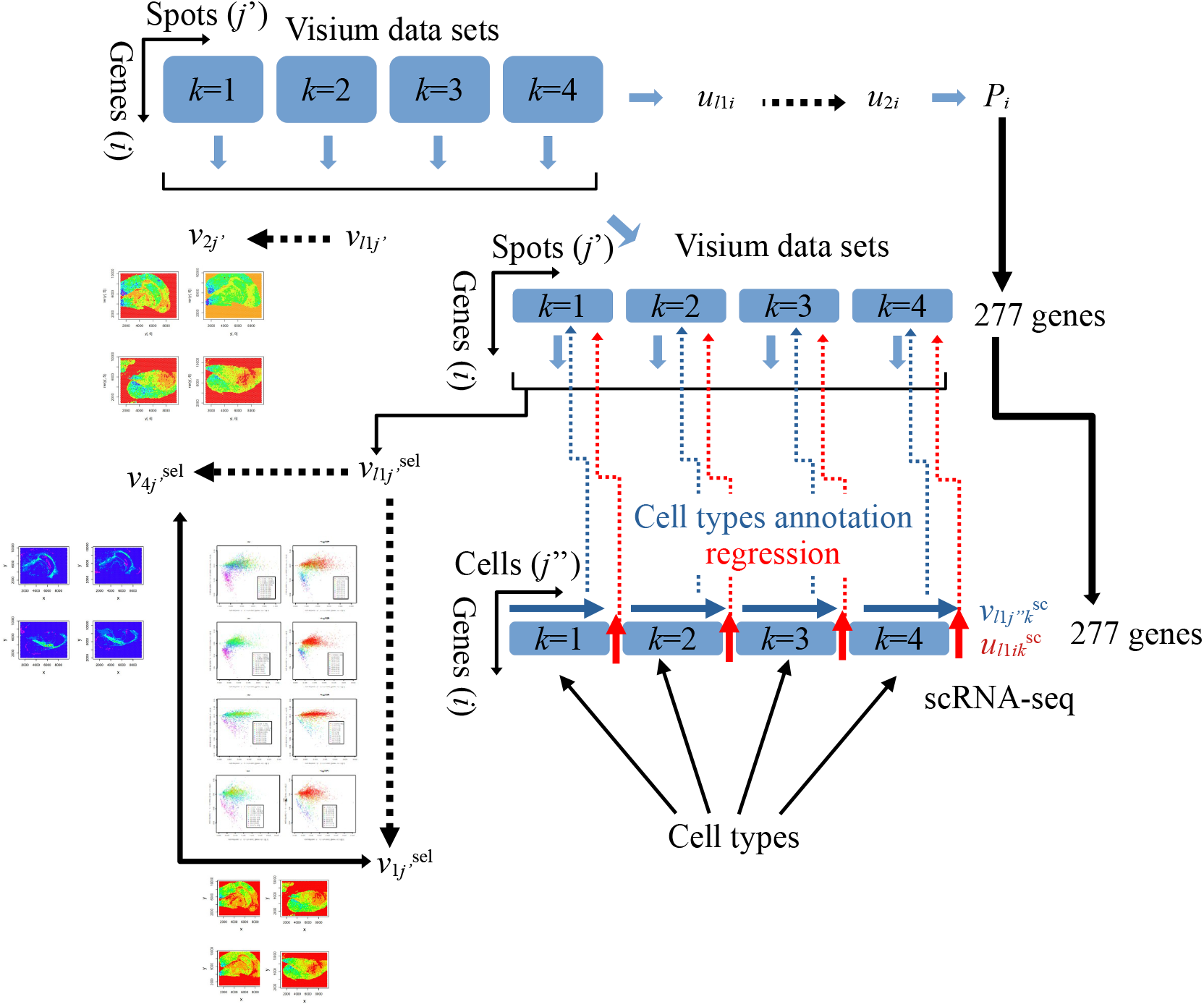
First, we integrate four Visium datasets and get 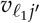 to see if it is consistent with the spatial coordinate. Then, as we found that *v*_2*j′*_ is consistent with the spatial coordinate, the corresponding *u*_2*i*_ is used to select 277 genes with adjusted *P*-values less than 0.01. Using 277 genes, 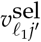 is computed and 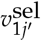 is still consistent with the spatial coordinate. Regression analysis is then performed wherein *x*_*ij′*_ is fitted with 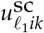 and the spot-associated significant *P*-values are selected. *P*-values are especially related to 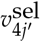. Finally, cell types of spots associated with significant adjusted *P*-values are inferred based on 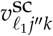.

**Figure 2.**
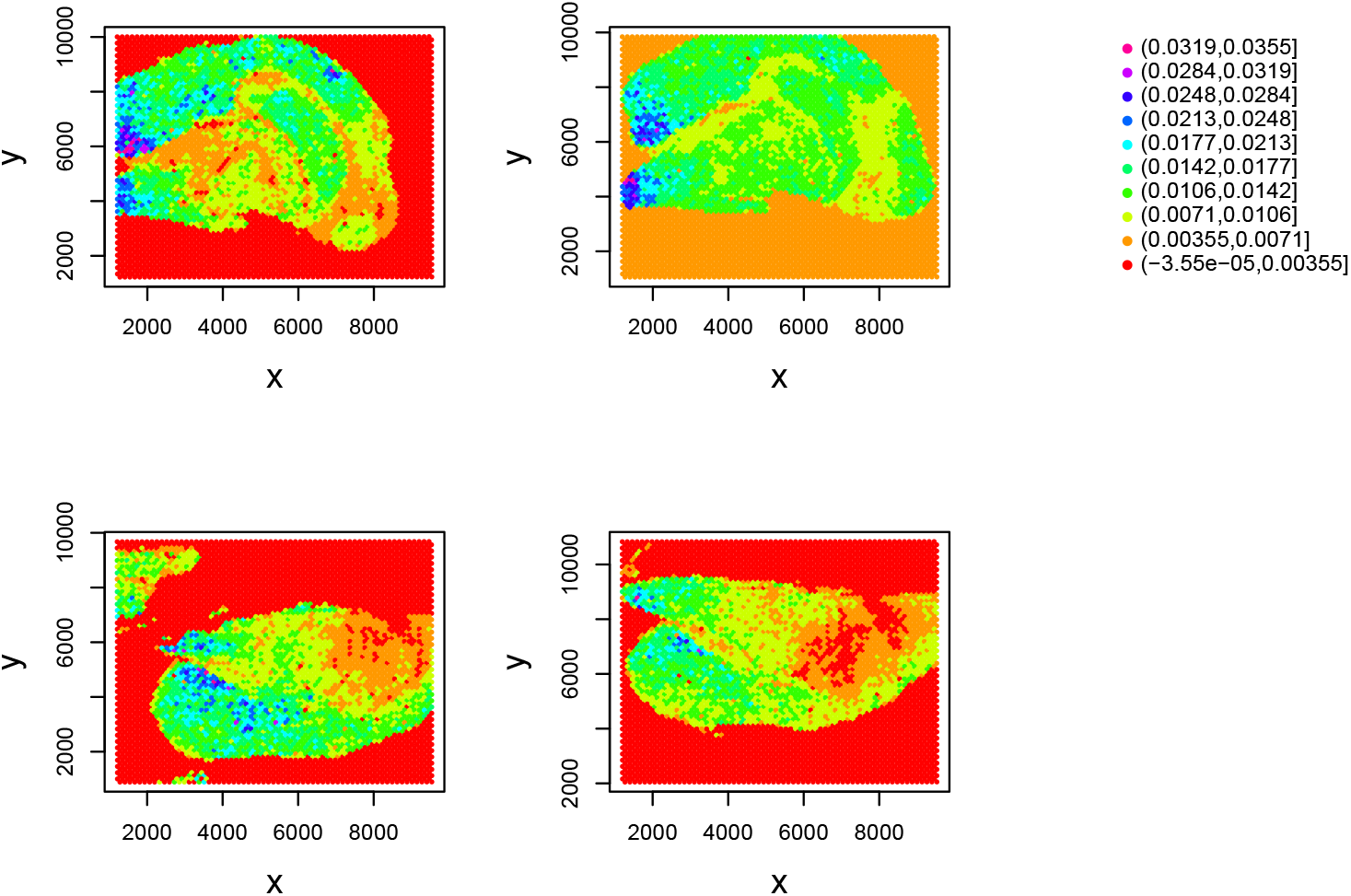
*v*_2*j′*_ attributed to the location of the *j′*th spot. Top left: *k* = 1, 1 ≤ *j′* ≤ 4992. Top right: *k* = 2, 4993 ≤ *j′* ≤ 9984. Bottom left: *k* = 3, 9985 ≤ *j′* ≤ 14976. Bottom right: *k* = 4, 14977 ≤ *j′* ≤ 19968.

### 2.6. Biological interpretation

Next, we would like to see if TD-based unsupervised FE applied to Visium can be used for biological interpretation, which is unclear because individual spots in Visium include multiple cells. Without annotations of individual spots, interpreting outcomes is difficult. One possible annotation to be attributed to individual spots is the cell fraction, i.e., deconvolution. Fortunately, as we have associated the single-cell gene expression profile, spots can be annotated based on single-cell RNA-seq. To do this, we need to annotate individual cells in single-cell RNA-seq.

Next we decided to employ 277 genes as biomarker genes to relate the gene expression of individual spots to that of single-cell RNA-seq. We computed SVD, 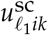, of single-cell RNA-seq using only 277 genes. Linear regression

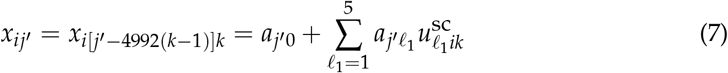

was performed to evaluate if 277 gene expression profiles in the Visium dataset can be represented by SVD computed from single-cell RNA-seq where 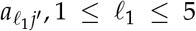 are regression coefficients. Figure 3 shows the spatial distribution of 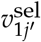 and the Spearman correlation coefficient between *x*_*ij′*_ and 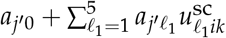, i.e., the left and right hand sides of eq. (7). The reason why not 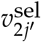 but 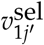 is because 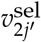 does not exhibit consistency with the spatial coordinate whereas 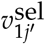 does. Not only is 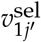 still consistent with the spatial coordinate, *x*_*ij′*_ is well computed from 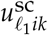. Table 2 lists the number of spots associated significant correlation between *x*_*ij′*_ and 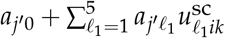. Obviously, gene expression of most spots has a significant correlation with SVD computed from single RNA-seq.

**Table 2.** the number of spots associated significant correlation.

| $k$ | Adjusted $P$-values | |
| --- | --- | --- |
| | $> 0.05$ | $\leq 0.05$ |
| 1 | 971 | 2076 |
| 2 | 1271 | 1769 |
| 3 | 1261 | 1349 |
| 4 | 1230 | 1351 |

**Figure 3.**
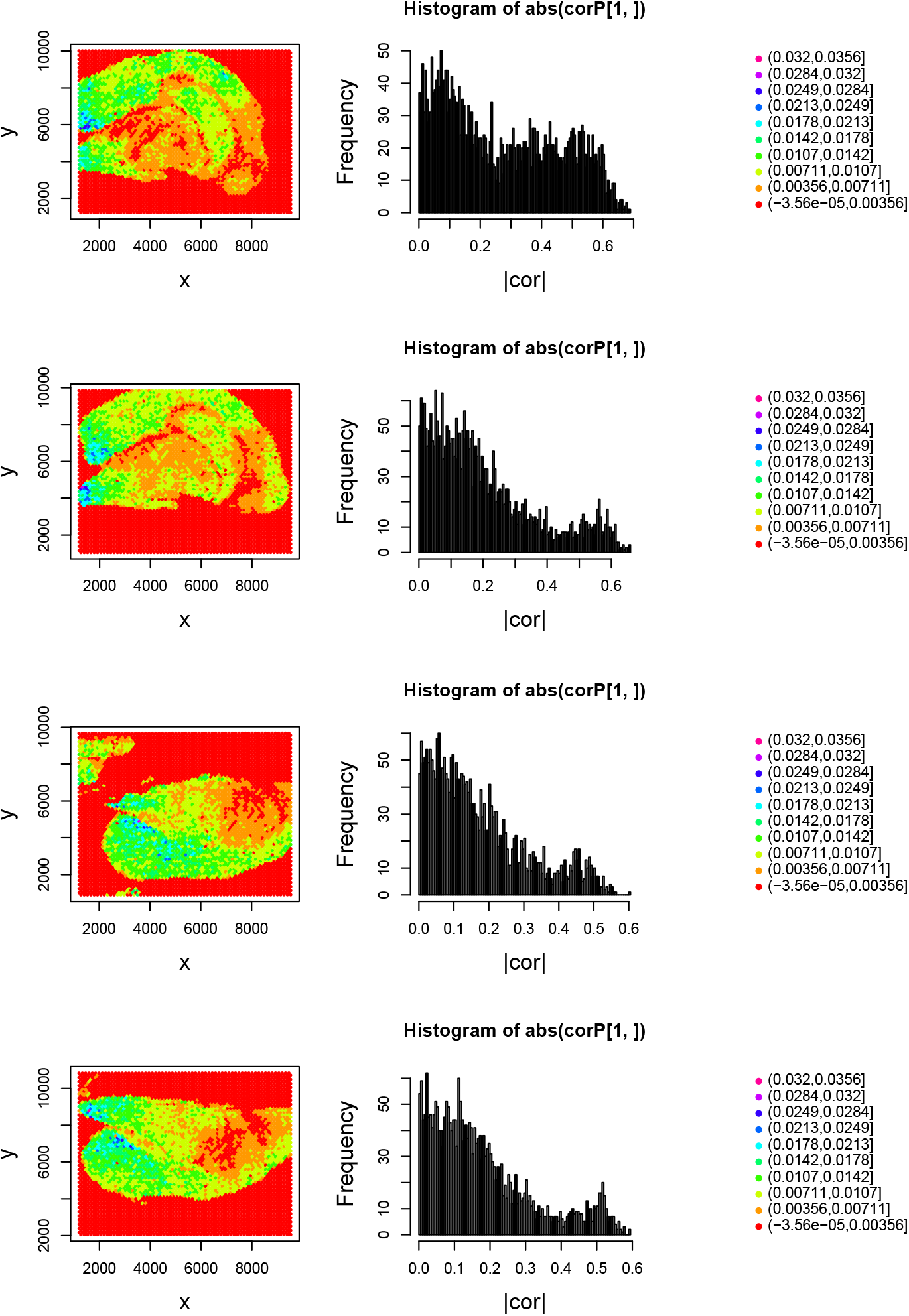
Left column: 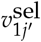 attributed to the location of the *j′*th spot. Right column: Histogram of the Spearman correlation coefficient between *x*_*ij′*_ and 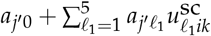. The first row: *k* = 1, 1 ≤ *j*^*′*^ ≤ 4992. The second row: *k* = 2, 4993 ≤ *j′* ≤ 9984. The third row: *k* = 3, 9985 ≤ *j′* ≤ 14976. The fourth row: *k* = 4, 14977 ≤ *j′* ≤ 19968.

Third, we would like to know the condition wherein spots are associated with significant correlation. For this purpose, we compute the Spearman correlation coefficient between 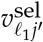 and negative signed logarithmic *P*-values of linear regression. We find that 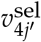 has the largest correlation coefficient (Fig. 4). Figure 5 shows the scatter plot between 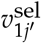 and 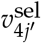 where colors correspond to the absolute Spearman correlation coefficient between *x*_*ij′*_ and 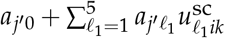 or negative-signed logarithmic *P*-values associated with the Spearman correlation coefficient. Obviously, the negatively larger 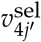 is associated with more significant correlation. To determine whether 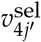 is still coincident with the spatial coordinate, we display 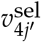 on spots (Fig. 6). Obviously, 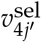 is still coincident with spatial coordinates.

**Figure 4.**
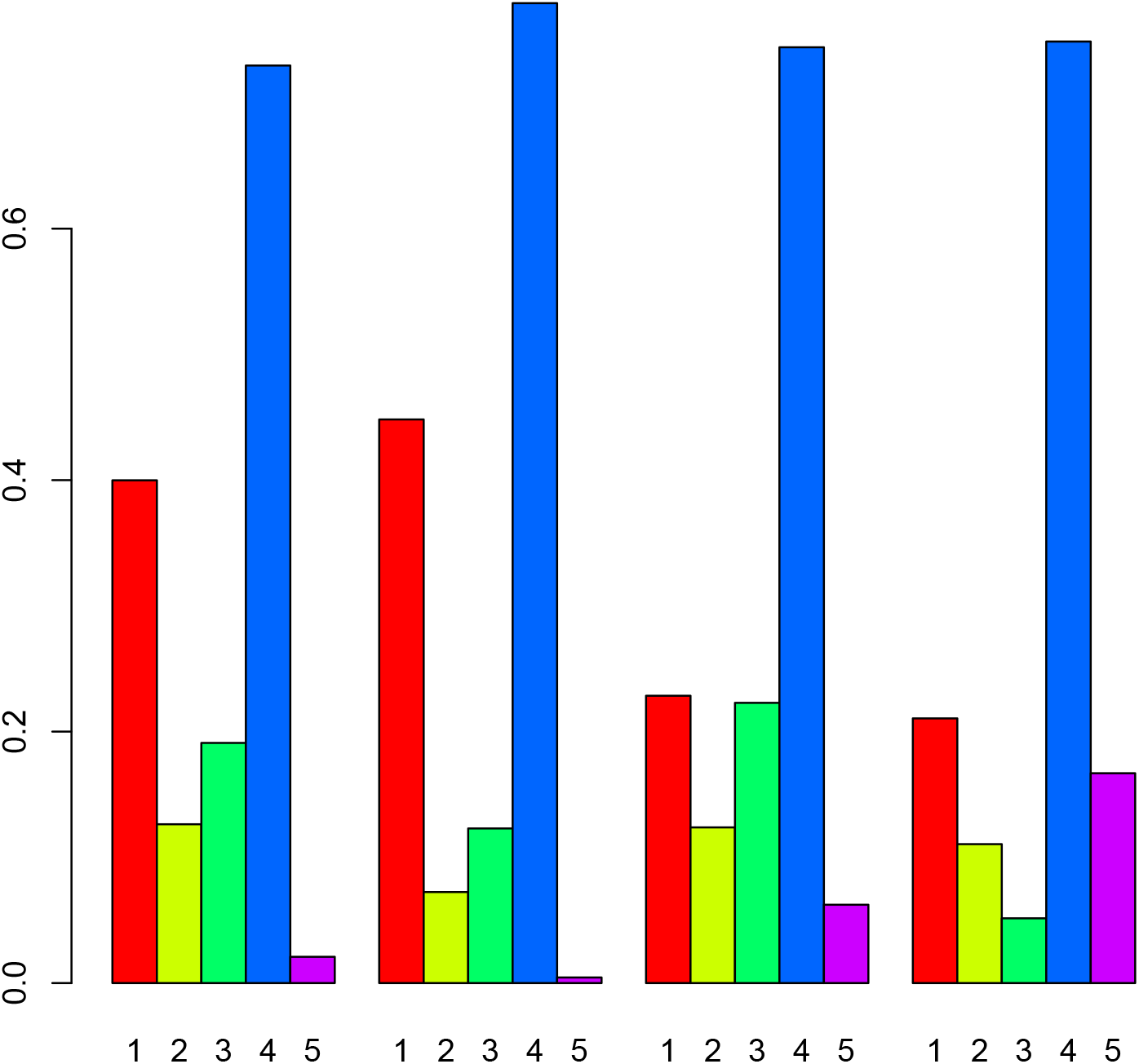
Absolute Spearman correlation coefficient between 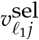 and negative-signed logarithmic *P*-values of linear regression, eq. (7). Numbers below horizontal axis correspond to *ℓ*_1_. From the left to right, *k* = 1, 2, 3, 4.

**Figure 5.**
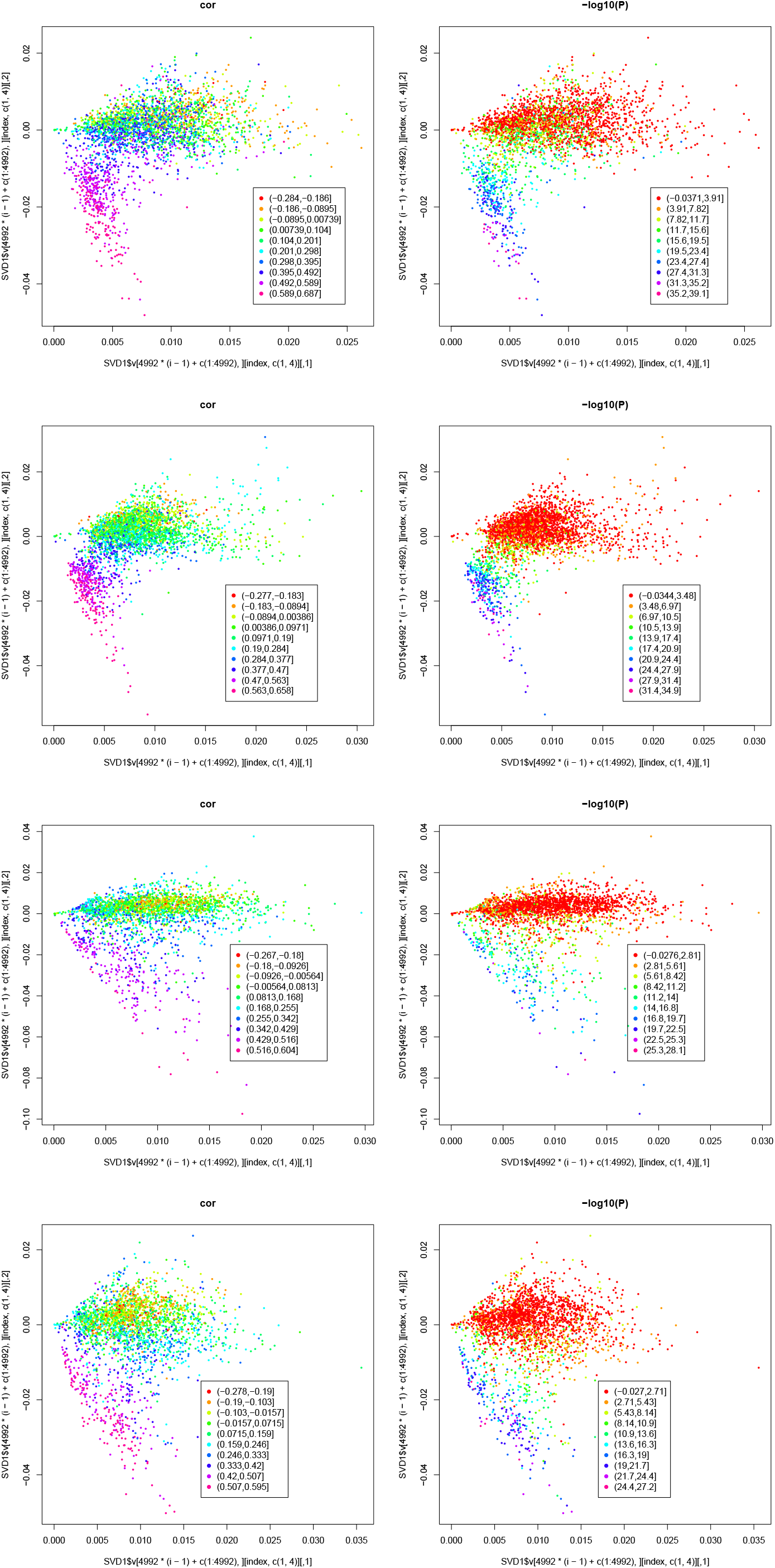
Scatter plot between 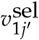 (horizontal axis) and 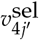 (vertical axis) where colors in the left column correspond to the absolute Spearman correlation coefficient between *x*_*ij′*_ and 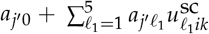 and those in the right column are negative-signed logarithmic *P*-values associated with the Spearman correlation coefficient. The first row: *k* = 1, 1 ≤ *j′* ≤ 4992. The second row: *k* = 2, 4993 ≤ *j′* ≤ 9984. The third row: *k* = 3, 9985 ≤ *j′* ≤ 14976. The fourth row: *k* = 4, 14977 ≤ *j′* ≤ 19968.

**Figure 6.**
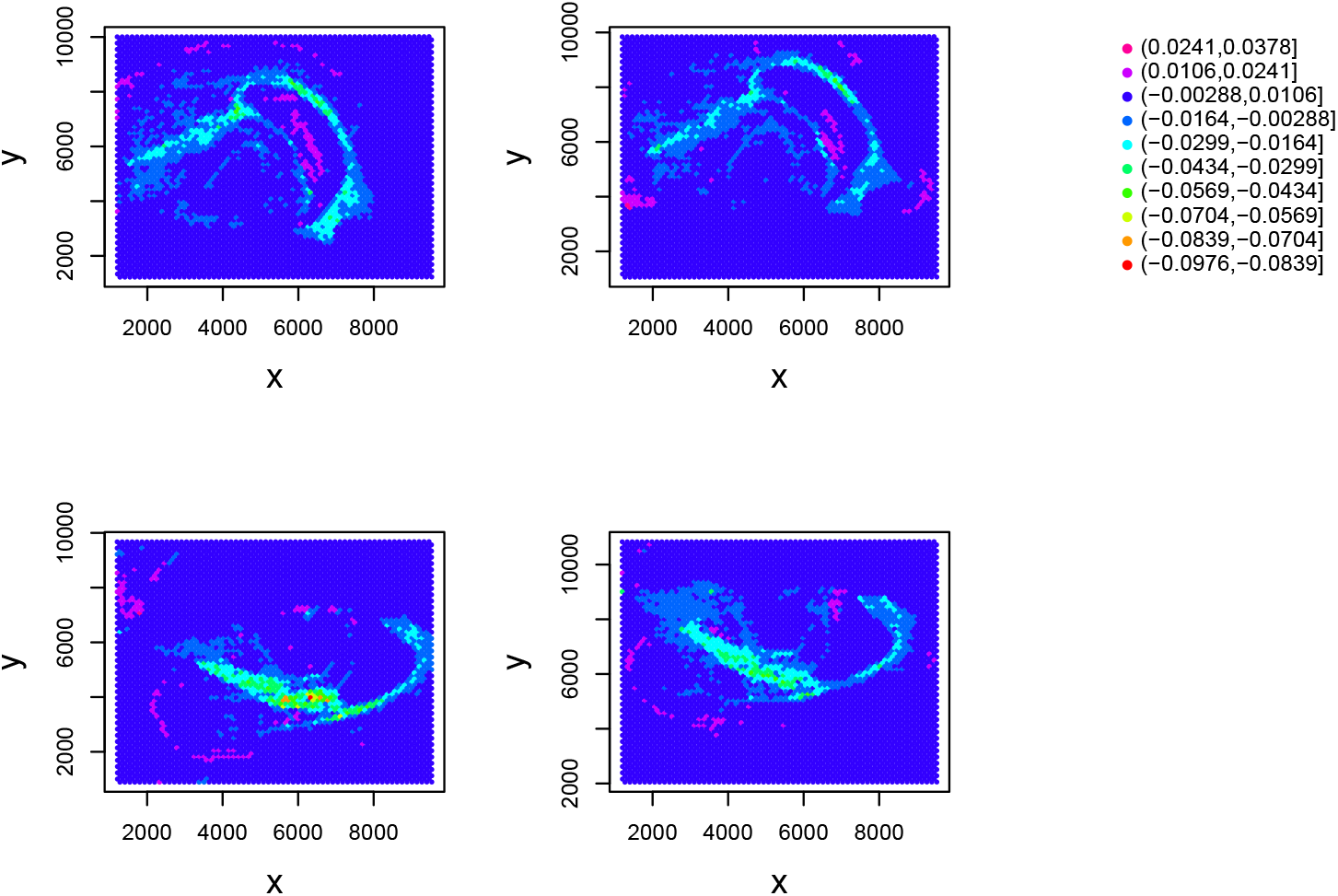
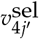 attributed to the location of the *j′*th spot. Top left: *k* = 1, 1 ≤ *j′* ≤ 4992. Top right: *k* = 2, 4993 ≤ *j′* ≤ 9984. Bottom left: *k* = 3, 9985 ≤ *j′* ≤ 14976. Bottom right: *k* = 4, 14977 ≤ *j′* ≤ 19968.

Then the fraction of the *s*th cell type, *c*_*j′* *s*_, of the *j′*th spot is computed as

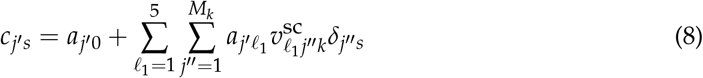

where *δ*_*j*′′*s*_ takes 1 only when the *j′′*th cell in single-cell RNA-seq belongs to the *s*th cell type otherwise 0. *M*_*k*_ is the total number of single cells in the *k*th single-cell RNA-seq. 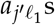 s are those computed in eq. (7). *c*_*j′* *s*_ was computed only for spots associated with adjusted *P*-values less than 0.05 in Table 2. This is the result shown in Figs. 20, 21, and 22.

## 3. Results

### 3.1. Failure of existing methods

First, we would like to demonstrate how poorly deconvolution is performed by some of conventional methods when reference single-cell gene expression includes multiple minor cell types (Table 1). As observed, neurons, micoroglia, and oligodendrocytes form the majority cell, consistent with the conclusion of the original paper [15]. Thus, the primary test of the following deconvolution methods is whether the majority cell types inferred by several methods include these three cell types. Below, deconvolution was performed using four conventional methods: RCTD [4], SPOTlight [5], SpaCET [6] and cell2location [7], which are briefly introduced in the Introduction, as well as with TD-based unsupervised FE. Figure 7 illustrates this analysis.

**Figure 7.**
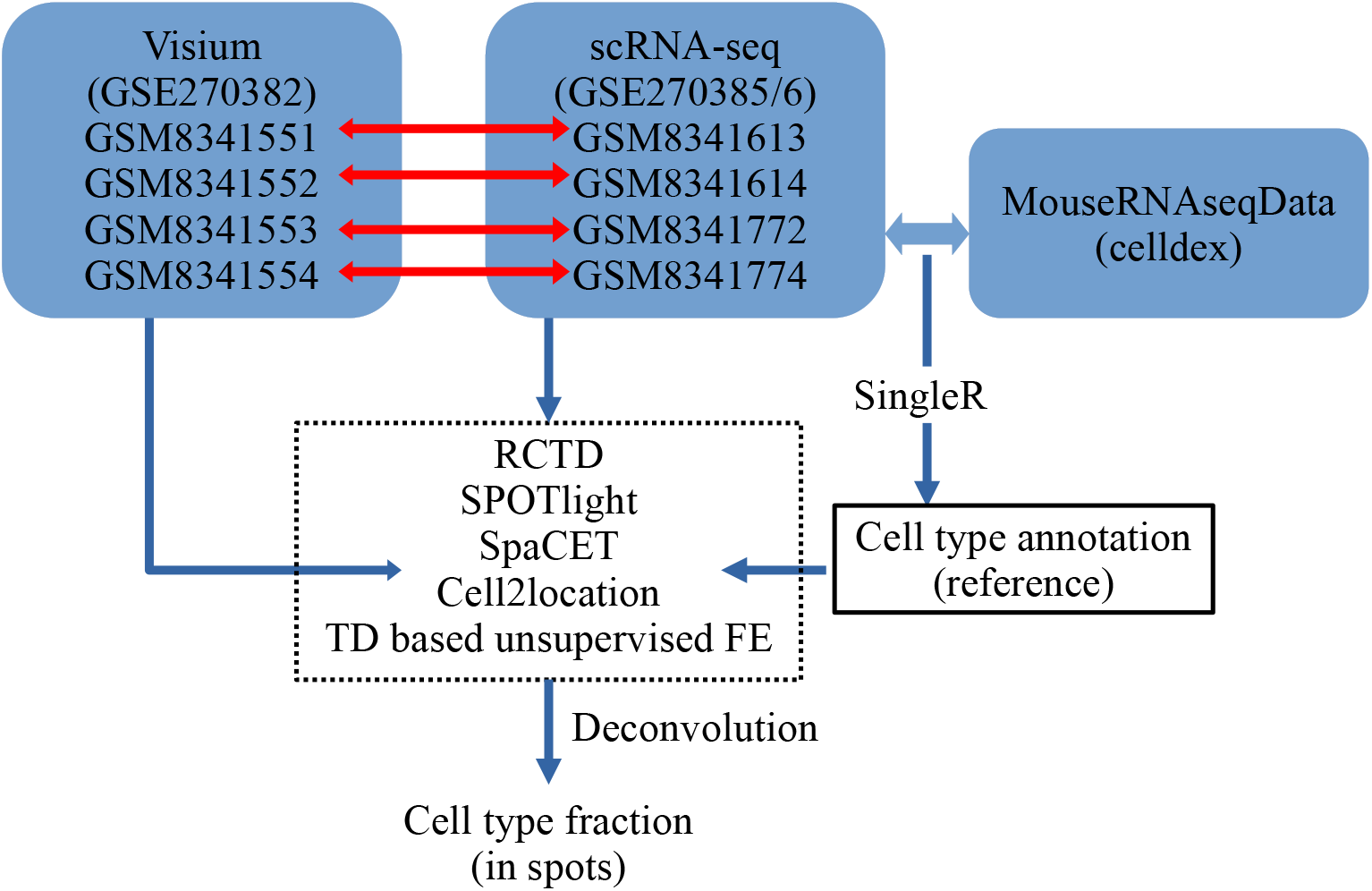
Flow chart of the analyses used in this study. Single-cell RNA-seq profiles are annotated using SingleR with the reference of MouseRNAseqData from celldex. Annotated cell types and single-cell RNA-seq are used to deconvolute Visium datasets with RCTD, SPOTlight, SpaCET, cell2location, and TD-based unsupervised FE. Horizontal red arrows show the correspondence between Visium datasets and reference single-cell RNA-seqs.

#### 3.1.1. RCTD

To determine whether conventional methods can perform deconvolution successfully, even when the reference single-cell gene expression profile includes minor cell types, as shown in Table 1, we first tested RCTD [4], a SOTA that aims for deconvolution (Fig. 8). Because cell deconvolution attributes multiple cell types to numerous individual spots, showing all of them in a visible way is difficult. Thus, we first plotted only the first 100 spots in the individual analyses and investigated more global features later in other visualizations. The primary problem is that RCTD detects many astrocytes that were reported to be absent in the original paper [15]. Thus, obviously, RCTD is unsuccessful. The reason of this failure is that RCTD cannot consider cell populations in the reference single-cell gene expression. The model used in RCTD is as follows. Suppose that there are *M* spots, where the expression of *N* genes is measured. Then *x*_*ijk*_ is modeled by RCTD as

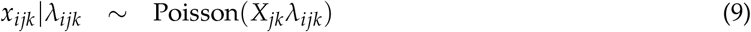

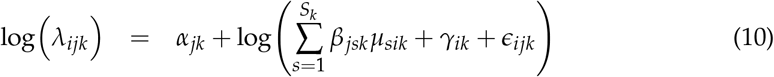

where *X*_*jk*_ is the total transcript count in the *j*th spot of the *k*th Visium dataset, *S*_*k*_ is the number of cell types present in the *k*th Visium dataset, *α*_*jk*_ is a fixed spot-specific effect, *µ*_*sik*_ is the mean gene expression profile for cell type *s* and gene *i, β*_*jsk*_ is the proportion of the contribution of cell type *s* to spot *j, γ*_*ik*_ is a gene-specific platform random effect, *ϵ*_*ijk*_ is a random effect to account for other sources of variation, such as spatial effects. As RCTD does not consider cell population in the reference, it cannot infer cell populations in individual spots of Visium data based on the cell populations in the reference. This is why RCTD wrongly counts too many (not existing) astrocyte populations in individual spots even though astrocytes are actually missing.

**Figure 8.**
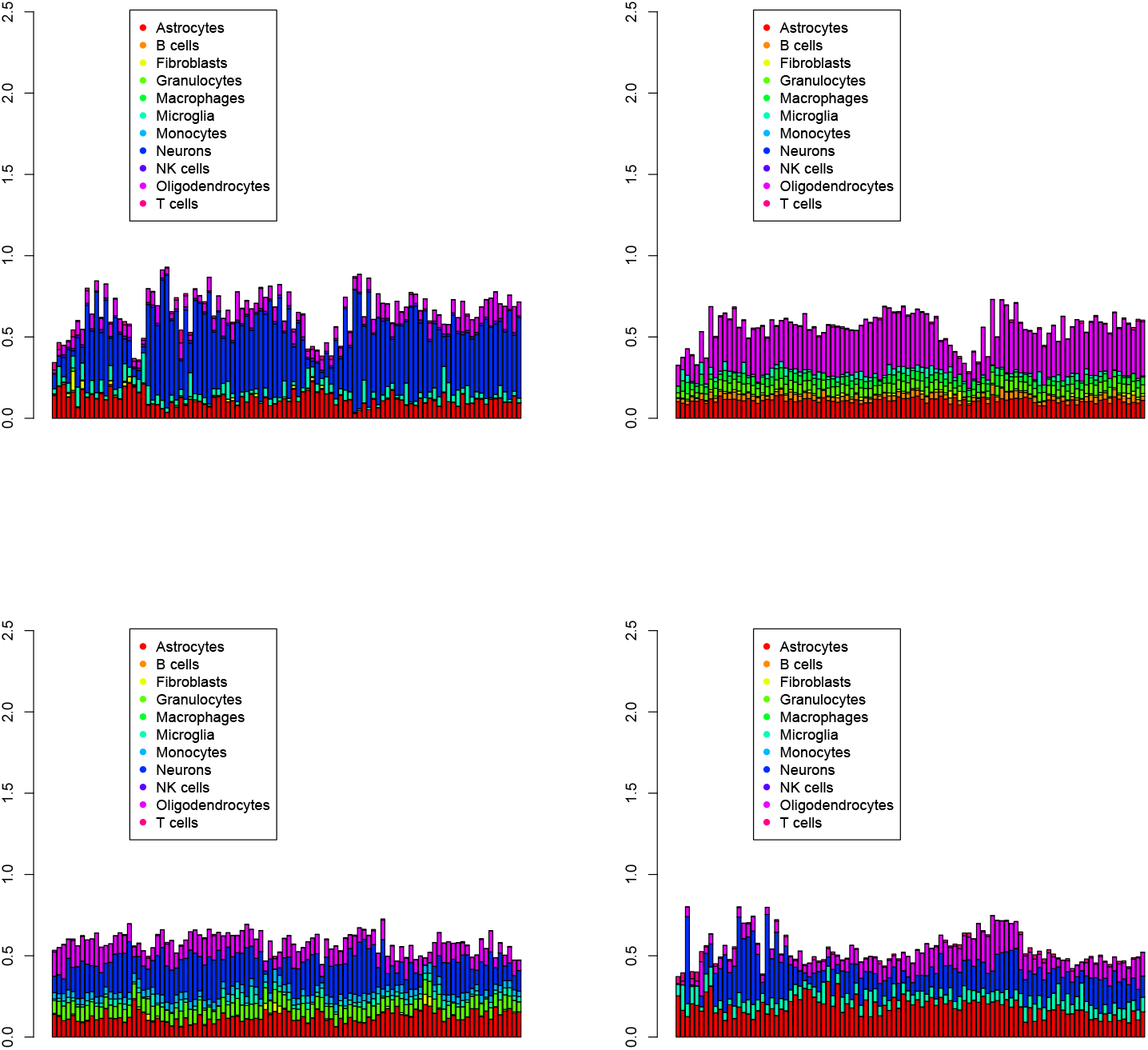
The first 100 cell populations for four Visium datasets (1 ≤ *k* ≤ 4) provided by RCTD. Top left: *k* = 1, top right: *k* = 2, bottom left: *k* = 3, bottom right *k* = 4.

#### 3.1.2. SPOTlight

Next we employed SPOTlight [5] (Fig. 9). As seen in Fig. 9, for *k* = 2, 3, the cell proportion is too heterogeneous to be true. This is possibly because of the marker gene identification step in SPOTlight that selects marker genes prior to deconvolution. SPOTlignt selects marker genes that are distinct among all cell types as much as possible; this results in an unintended emphasis toward minor cell types and unrealistic heterogenous cell proportions. Thus, SPOTlight is also unsuccessful.

**Figure 9.**
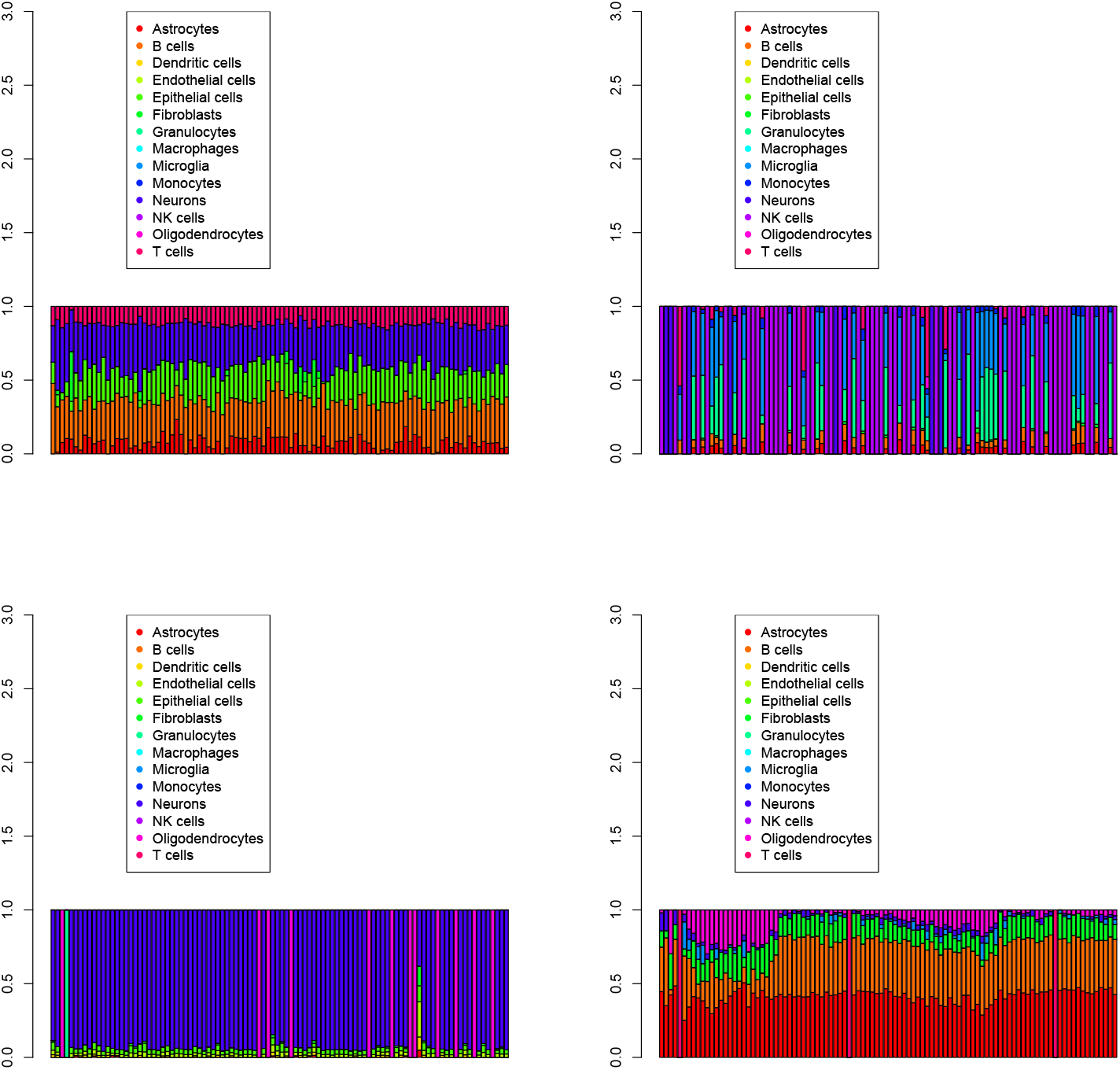
The first 100 cell populations for four Visium datasets (1 ≤ *k* ≤ 4) provided by SPOTlight. Top left: *k* = 1, top right: *k* = 2, bottom left: *k* = 3, bottom right *k* = 4.

#### 3.1.3. SpaCET

Although SPOTlight and RCTD are SOTA, because they are a bit old fashioned, we employ SpaCET [6] as a new alternative to RCTD and SPOTlight. Although SpaCET has been designed to process tumors, it can process gene expression other than that of tumors when reference-single-cell gene expression profiles associated with Visium data are available (Fig. 10). As many cell types other than neurons, micoroglia, and oligodendrocytes, which are the majority in reference datasets (see Table 1), are the majority in the four Visium datasets (e.g., astrocytes for *k* = 1, dendritic cells, endothelial cells, and NK cells for *k* = 3), and cell fractions are too distinct between individual Visium datasets, SpaCET is hardly successful. The reason why SpaCET fails is that it considers only correlation coefficients of gene expression between individual single cells in the reference and the spots in Visium that cannot consider the cell type fraction in reference single-cell profiles, as in the case of RCTD.

**Figure 10.**
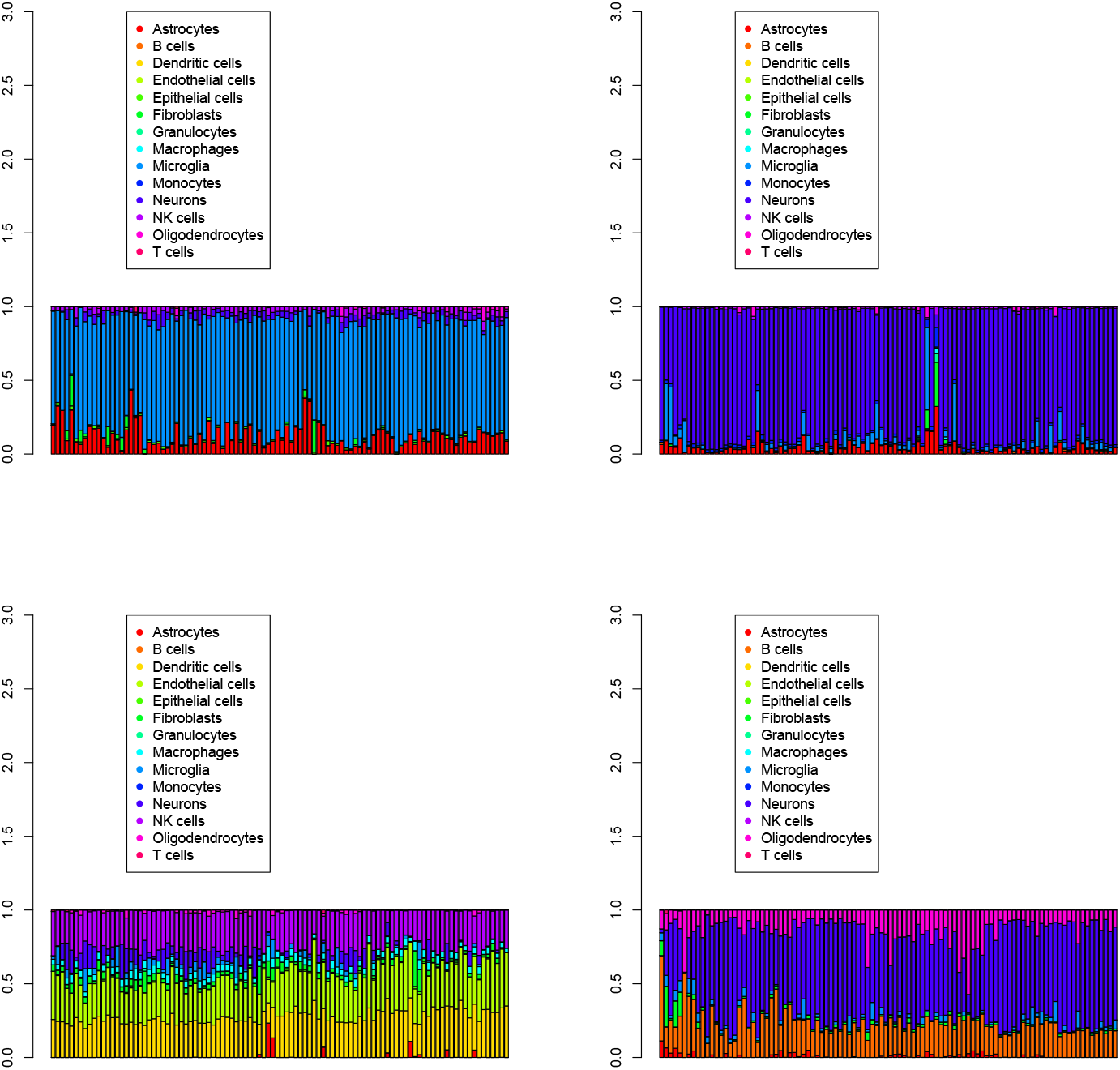
The first 100 cell populations for four Visium datasets (1 ≤ *k* ≤ 4) provided by SpaCET. Top left: *k* = 1, top right: *k* = 2, bottom left: *k* = 3, bottom right *k* = 4.

#### 3.1.4. cell2location

Finally, we test a more advanced Bayesian-based method called cell2location [7] (Fig. 11). Although the Bayesian method is expected to be more accurate and robust, it is very time-consuming because it must optimize the probability itself (and cell2location is greatly time-consuming). As none of the three majority cell types in the four Visium datasets were neurons, microglia, and oligodendrocytes, which were the majority in the reference datasets (see Table 1), cell2location is also unlikely to be successful despite a very long running time (it takes a few days, whereas the other three finish in less than at most a few tens of minutes).

**Figure 11.**
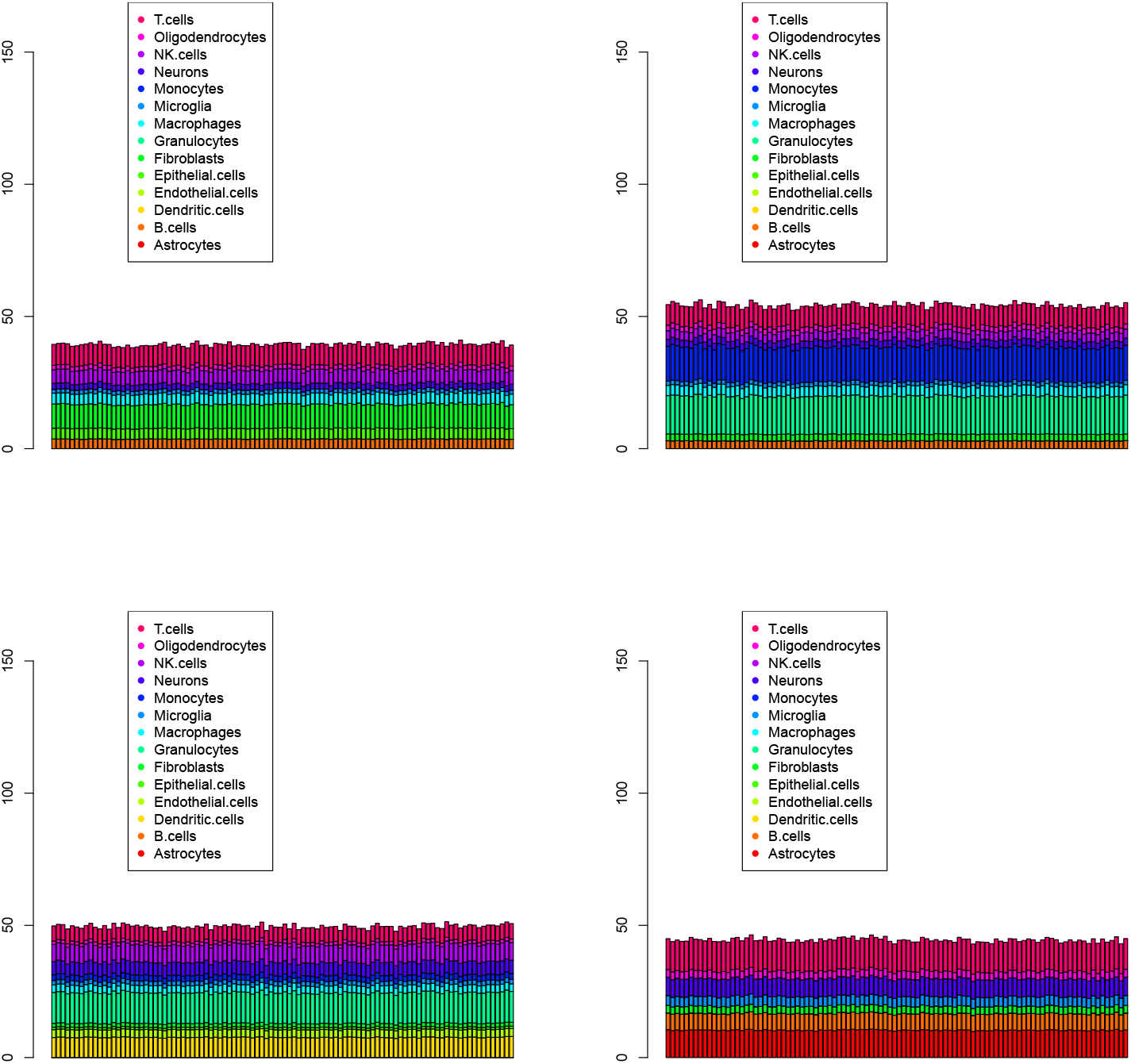
The first 100 cell population for four Visium datasets (1 ≤ *k* ≤ 4) provided by cell2location. Top left: *k* = 1, top right: *k* = 2, bottom left: *k* = 3, bottom right *k* = 4.

### 3.2. Over all cell type fraction

Although the visualization restricted to the first 100 spots is useful for quickly grasping how well the individual methods work, it lacks global information for all of the spots. To support this missing information, we present pie charts representing all fractions of the identified cell types over all of the spots.

#### 3.2.1. RCTD

As expected in Fig. 8, Astrocytes which should be missing (see Table 1) is one of primary cell types predicted by RCTD. In addition, one of the four pie charts had completely missing neurons, which should be the primary cell types (see Table 1), in Fig. 12. Clearly, RCTD failed to predict the correct cell-type fractions.

**Figure 12.**
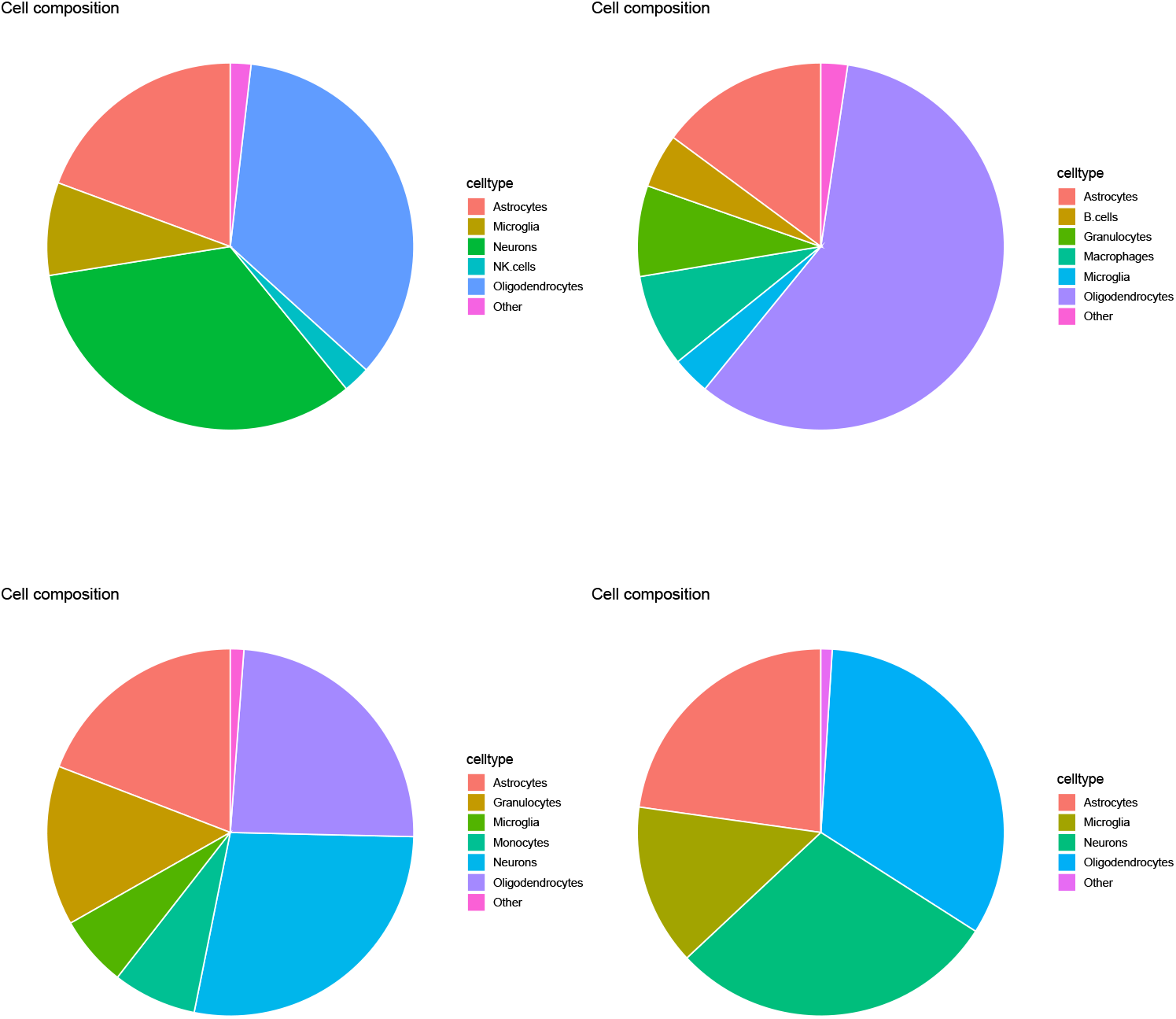
Pi charts of over all cell type fractions for four Visium datasets (1 ≤ *k* ≤ 4) provided by RCTD. Top left: *k* = 1, top right: *k* = 2, bottom left: *k* = 3, bottom right *k* = 4.

#### 3.2.2. SPOTlight

As expected in Fig. 10, four samples in Fig. 13 are too distinct from one another. For example, the neurons were dominant for *k* = 3 whereas Astrocytes that were missing were dominant for *k* = 4. Because such a distinct cell type fraction is unlikely to be true when we consider the cell type fraction in the reference (Table 1), SPOTlight failed to reproduce a reasonable cell type fraction.

**Figure 13.**
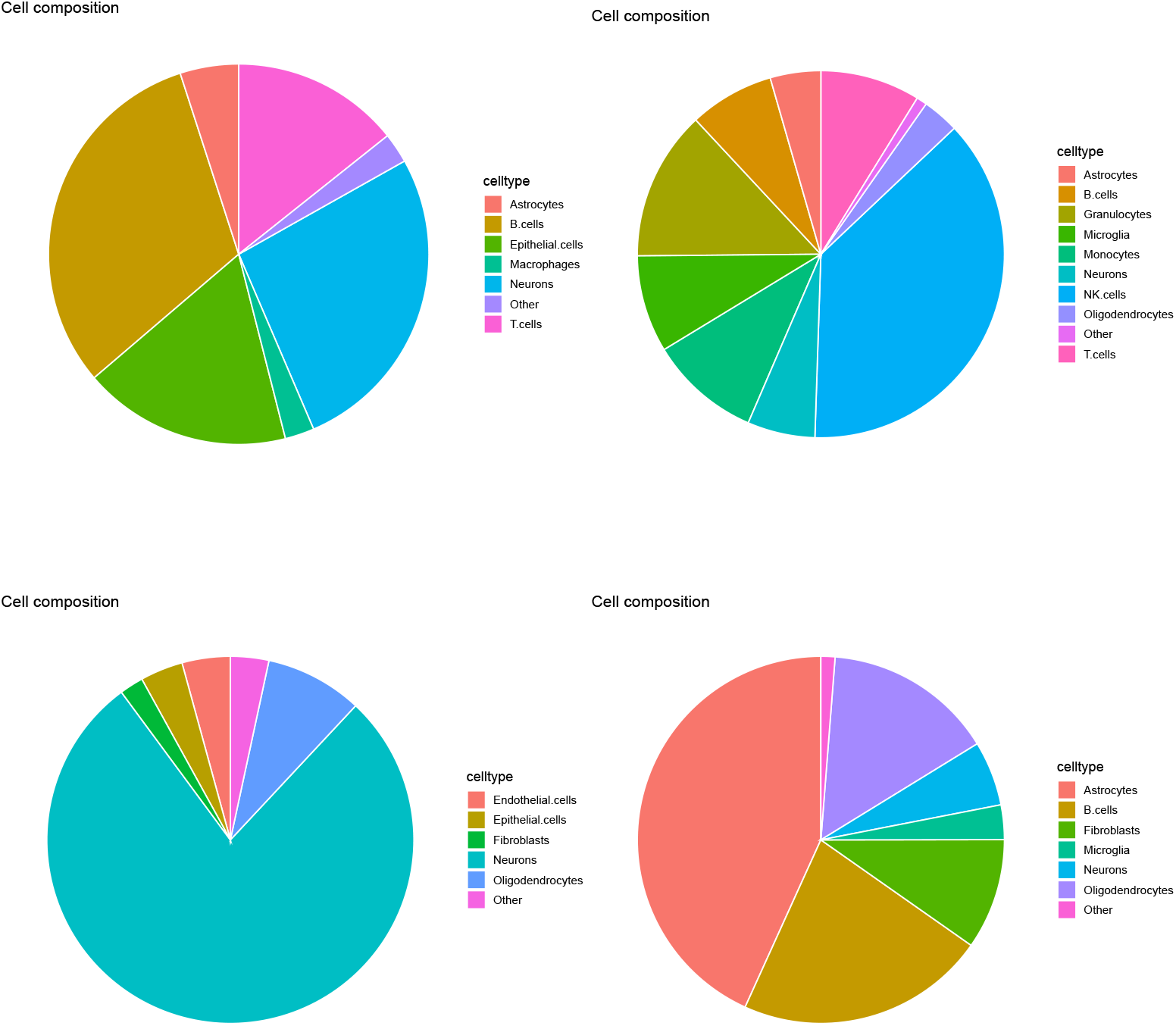
Pi charts of over all cell type fraction for four Visium datasets (1 ≤ *k* ≤ 4) provided by SPOTlight. Top left: *k* = 1, top right: *k* = 2, bottom left: *k* = 3, bottom right *k* = 4.

#### 3.2.3. SpaCET

Evaluation of SpaCET based on the Pi chart (Fig. 14) is the same as that based on Fig. 10; as many of cell types other than neurons, micoroglia, and oligodendrocytes, which were majorities in reference data sets (see Table 1), are majority in four Visium datasets (e.g., astrocytes for *k* = 1, dendritic cells, endothelial cells, and NK cells for *k* = 3) and cell fractions are too distinct between individual Visium datasets, SpaCET is hardly successful.

**Figure 14.**
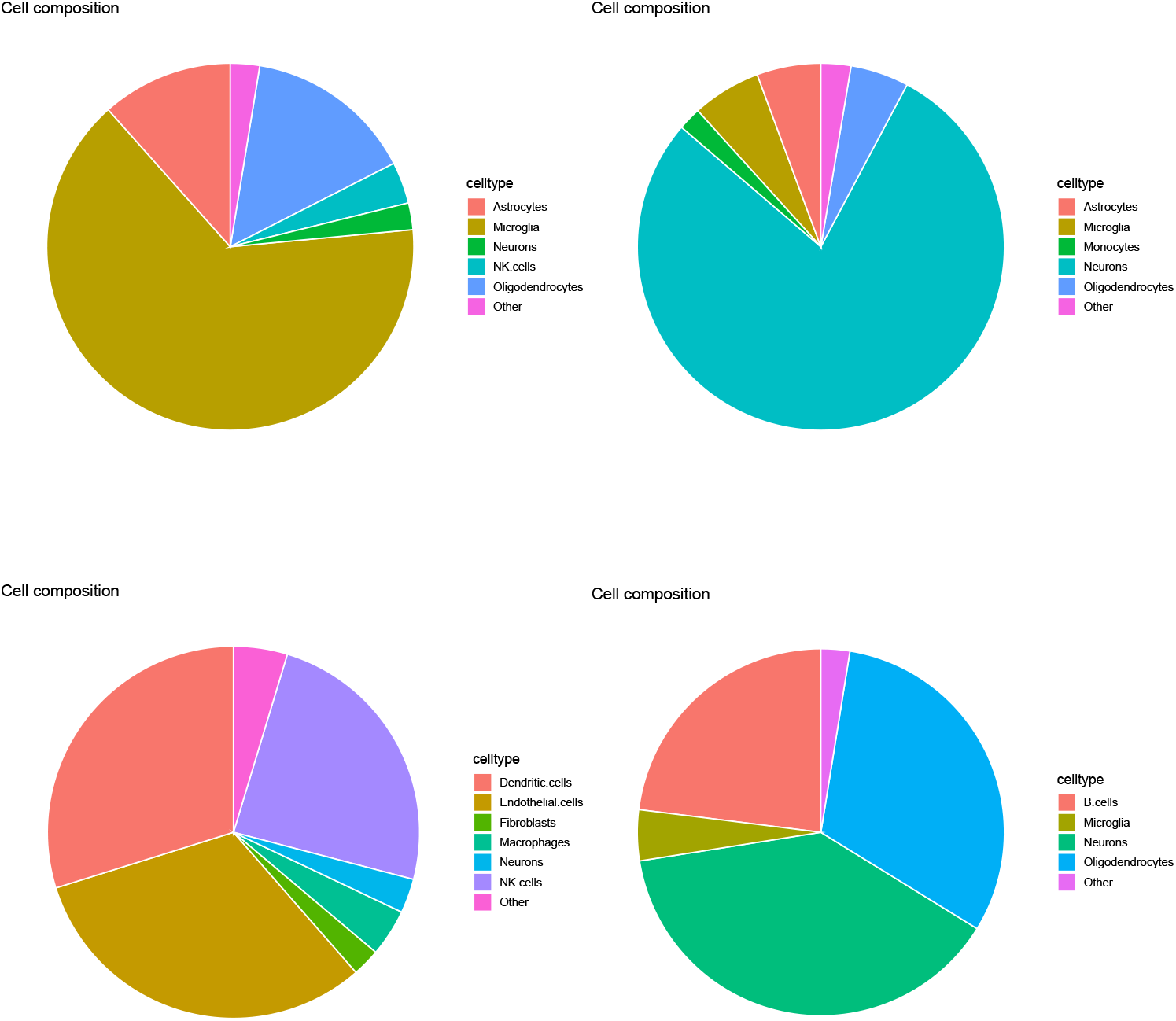
Pi charts of over all cell type fraction for four Visium datasets (1 ≤ *k* ≤ 4) provided by SpaCET. Top left: *k* = 1, top right: *k* = 2, bottom left: *k* = 3, bottom right: *k* = 4.

#### 3.2.4. cell2location

Evaluation of SpaCET based on the pi chart (Fig. 15) is the same as that based on Fig. 11; as none of three majority cell types in four Visium datasets are neurons, microglia, and oligodendrocytes, which were majorities in the reference data sets (see Table 1), cell2location is also unlikely to be successful.

**Figure 15.**
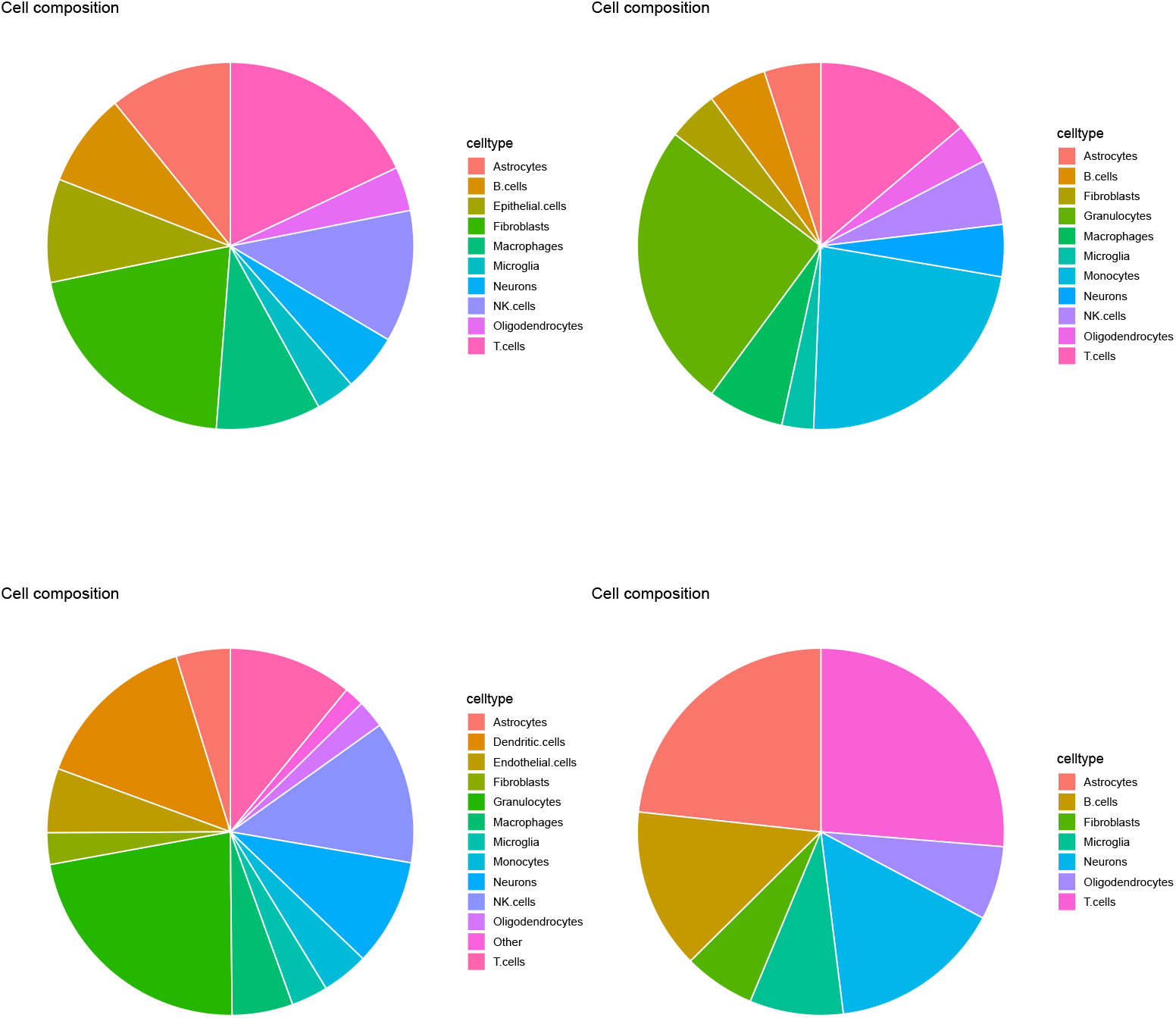
Pi charts of over all cell type fraction for four Visium datasets (1 ≤ *k* ≤ 4) provided by cell2location. Top left: *k* = 1, top right: *k* = 2, bottom left: *k* = 3, bottom right *k* = 4.

### 3.3. Spatial abundance of cell types

Although the overall cell type fraction shown in the pi charts is sufficient in order to show that these four methods are unsuccessful, we also show yet another evidence of unsuccess of four methods: spatial abundance of cell types. Here, we considered three major cell types in reference: microgiria, neurons, and oligodendrocytes (Figs. 16,17,18, and 19). Obviously, none of them had a reasonable spatial abundance, but all had a have random abundance of cell types. Clearly, they were unsuccessful.

**Figure 16.**
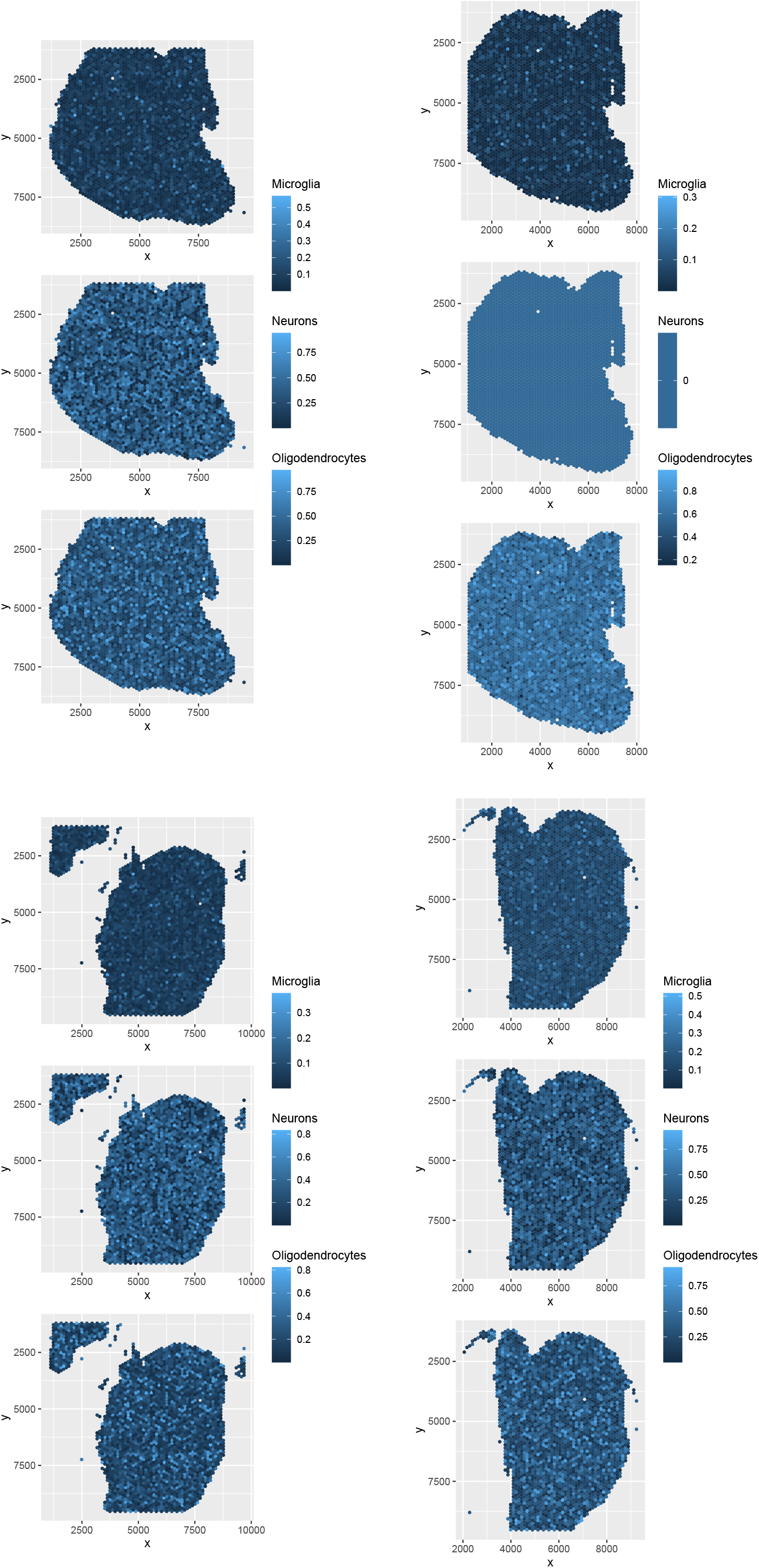
Spatial abundance of cell types for four Visium datasets (1 ≤ *k* ≤ 4) provided by RCTD. Top left: *k* = 1, top right: *k* = 2, bottom left: *k* = 3, bottom right: *k* = 4.

**Figure 17.**
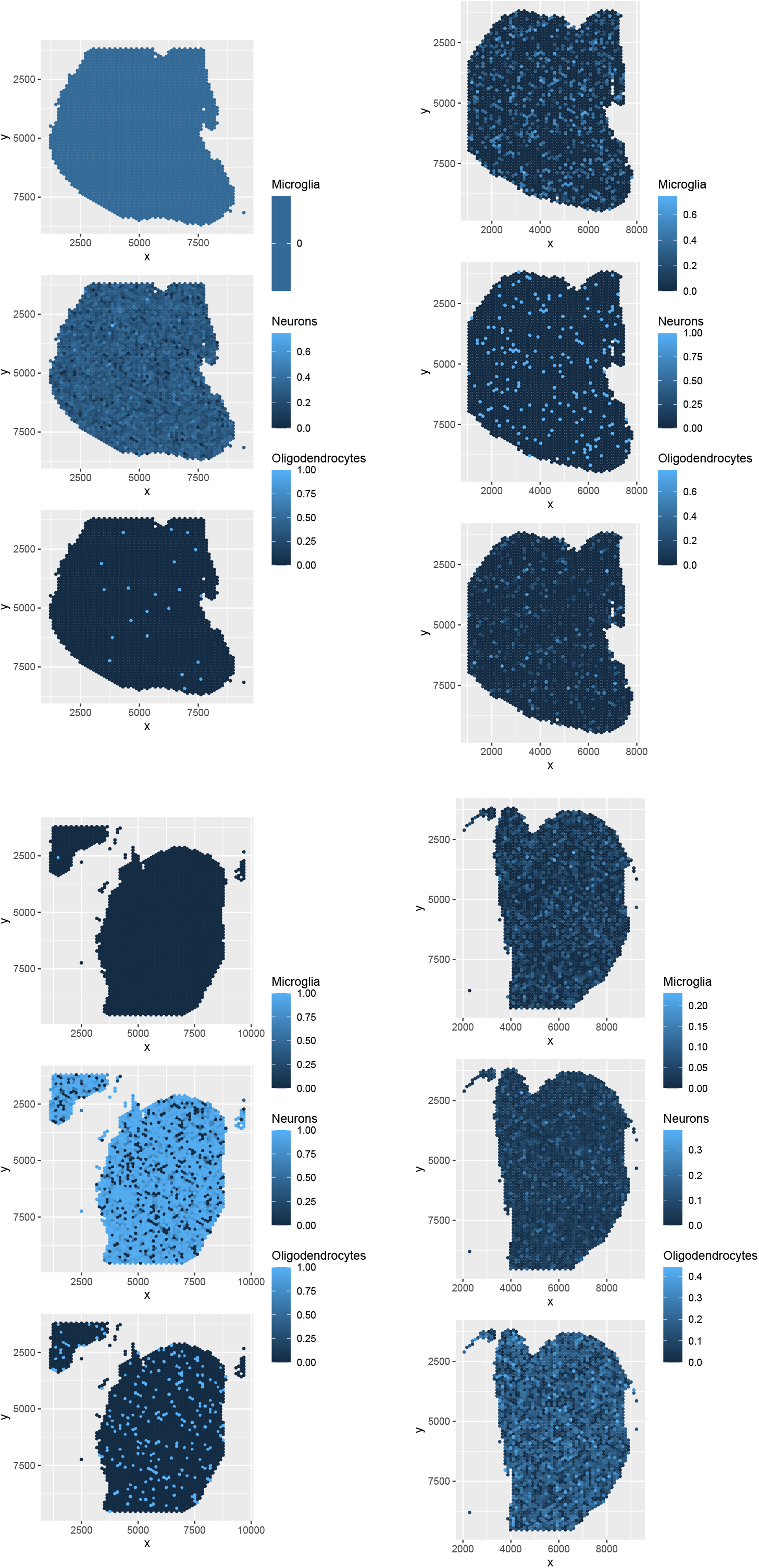
Spatial abundance of cell types for four Visium datasets (1 ≤ *k* ≤ 4) provided by SPOTlight. Top left: *k* = 1, top right: *k* = 2, bottom left: *k* = 3, bottom right: *k* = 4.

**Figure 18.**
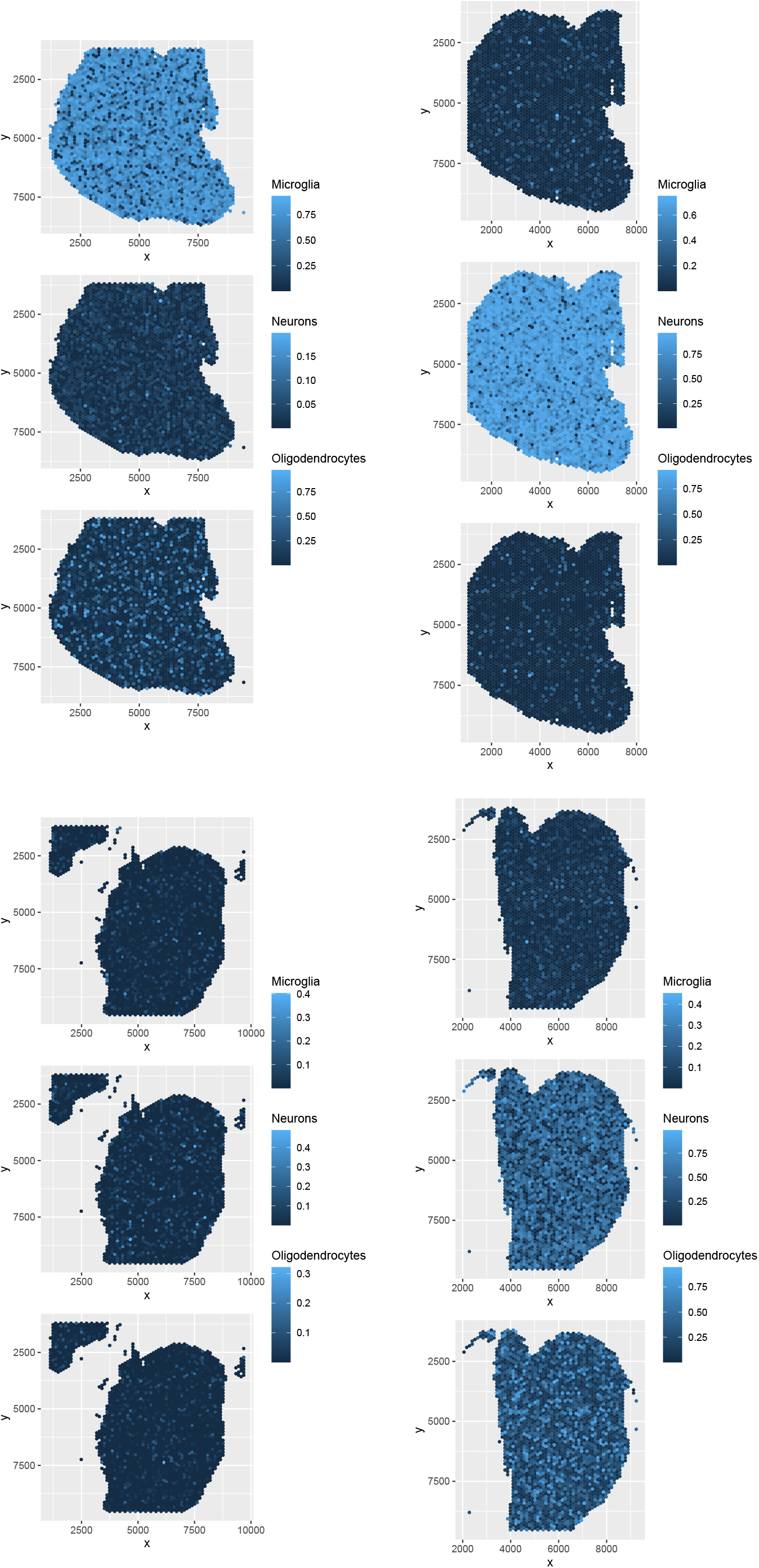
Spatial abundance of cell types for four Visium datasets (1 ≤ *k* ≤ 4) provided by SpaCET. Top left: *k* = 1, top right: *k* = 2, bottom left: *k* = 3, bottom right: *k* = 4.

**Figure 19.**
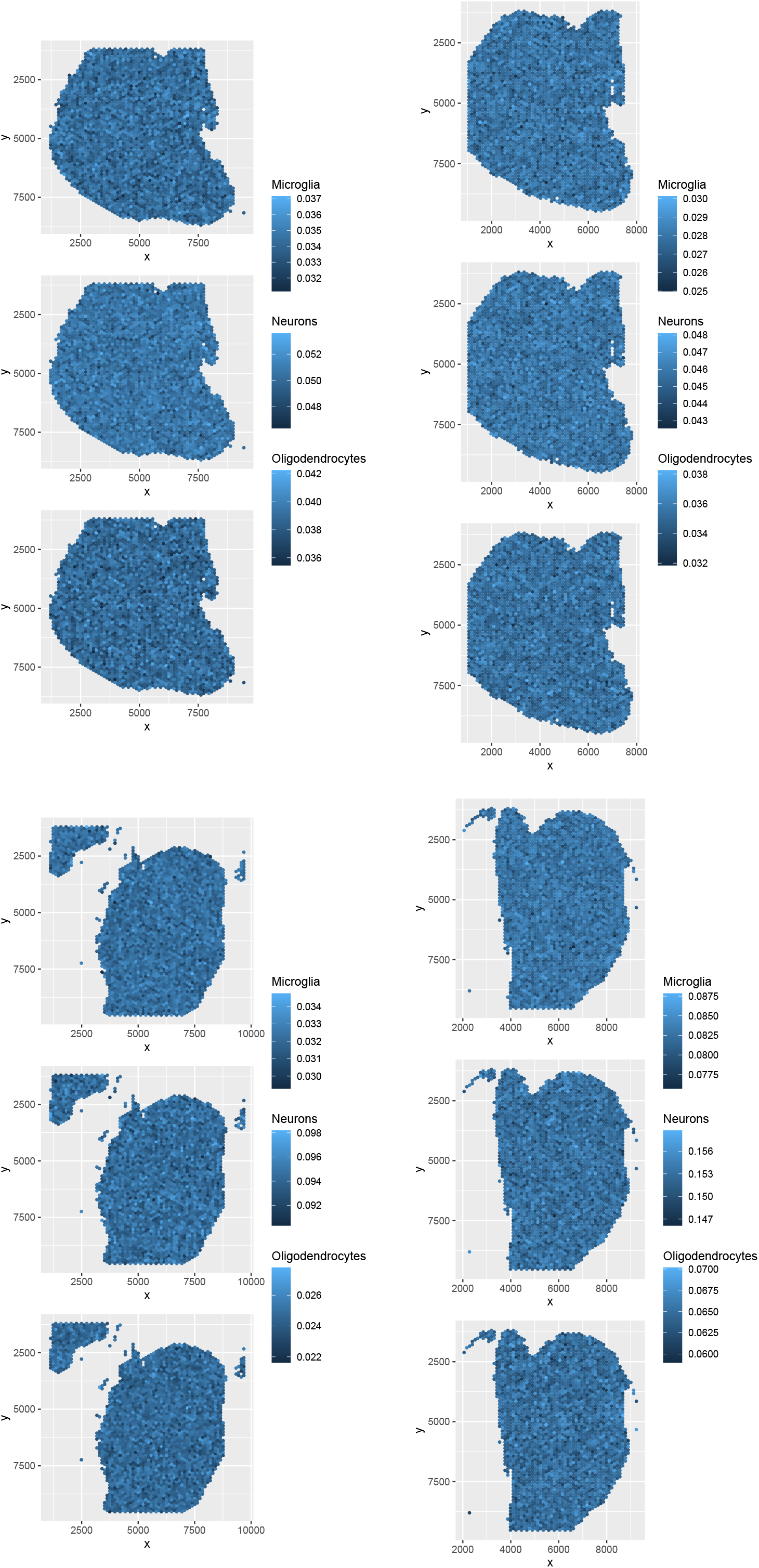
Spatial abundance of cell types for four Visium datasets (1 ≤ *k* ≤ 4) provided by cell2location. Top left: *k* = 1, top right: *k* = 2, bottom left: *k* = 3, bottom right *k* = 4.

### 3.4. TD based unsupervised FE applied to Visium

To determine whether TD-based unsupervised FE can outperform the above four methods, we applied TD-based unsupervised FE to Visium data. (To see how we obtained the results in this section, see the Methods section). Figure 20 shows *c*_*j′ s*_ (cell type profiles computed using TD-based unsupervised FE. See the following for further details) for 1 ≤ *k* ≤ 4: Clearly, only three cell types, “Microglia” “Neurons” and “Oligodendrocytes” and oligodendrocytes are dominant, which is consistent with the original study [15]. Thus, our inference of cell types is reasonable, and TD-based unsupervised FE is the only method to successfully process Visium datasets, even when the associated reference profiles include many minor cell types.

**Figure 20.**
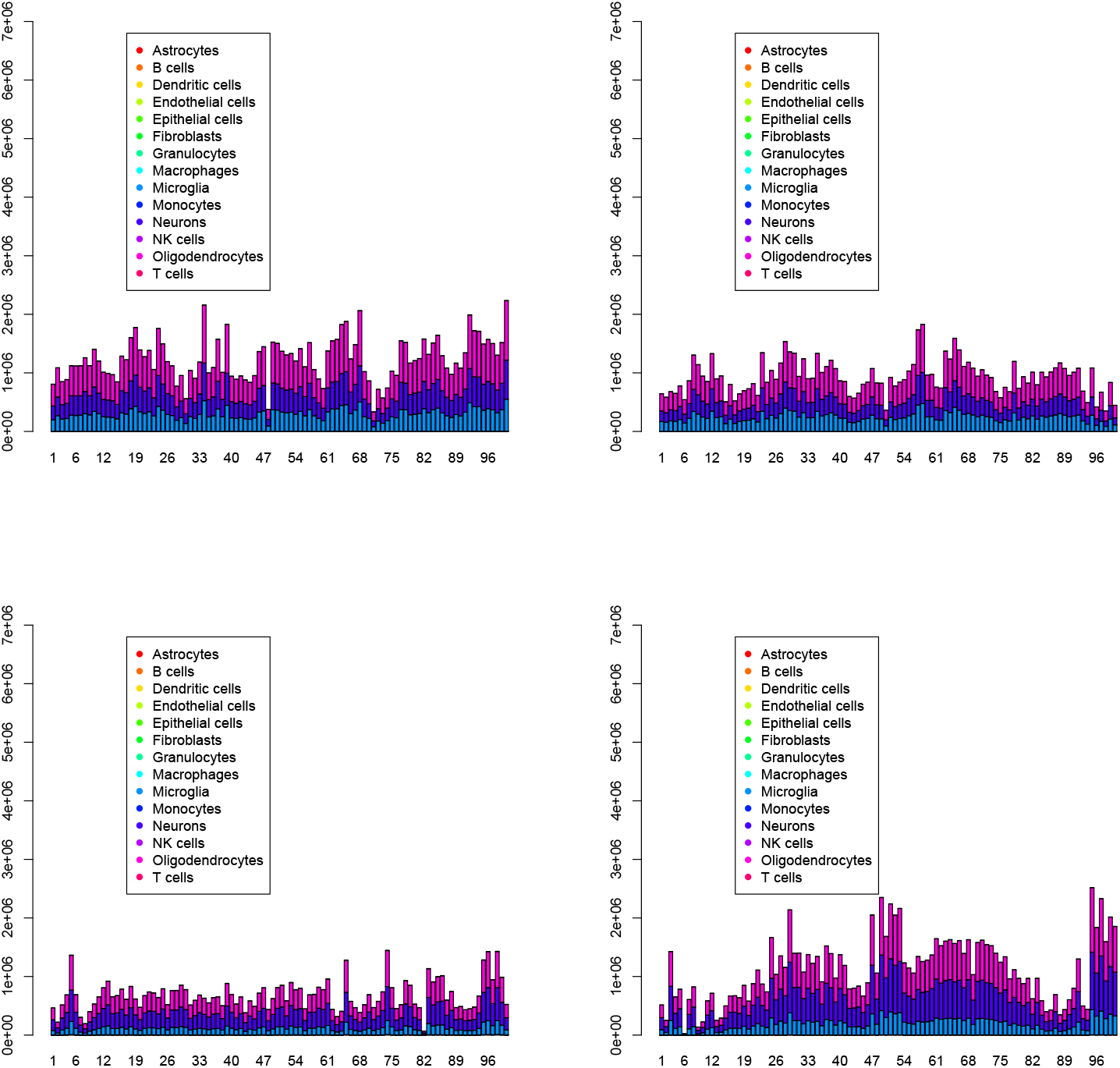
Pi charts of over all cell type fraction for four Visium datasets (1 ≤ *k* ≤ 4) by-TD based unsupervised FE. Top left: *k* = 1, top right: *k* = 2, bottom left: *k* = 3, bottom right *k* = 4.

We also show the pi chart and spatial abundance of cell types (Figs. 21 and 22). The reasons for the missing spots in Fig. 22 is because only spots whose gene expression is correlated with 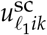 are plotted. This can be both advantageous and disadvantageous (see Discussion). The three major cell types are clearly identical to the reference (Table 1), and the spatial abundance is also reasonable. TD based unsupervised FE outperformed the other four methods.

**Figure 21.**
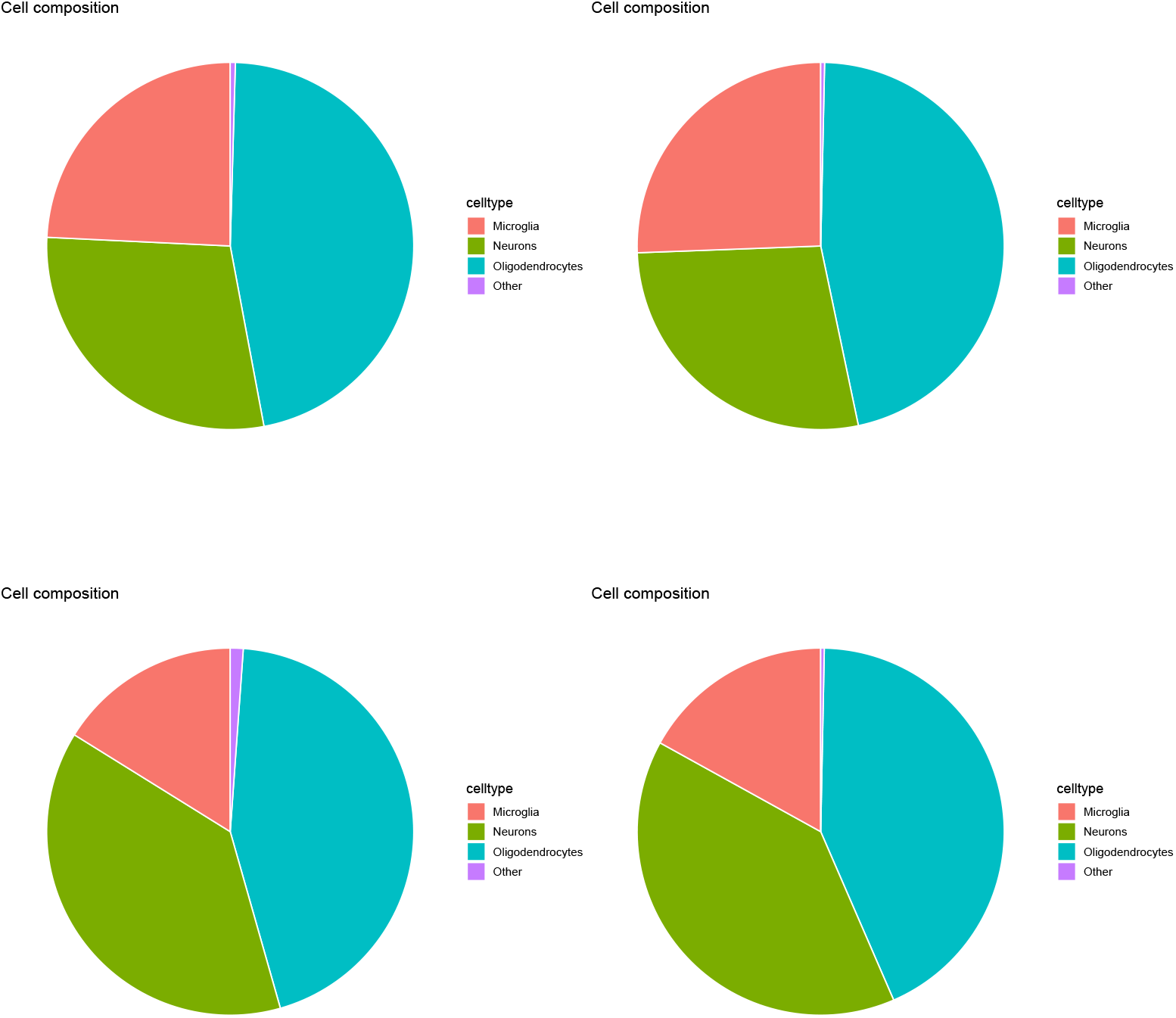
Pi charts of over all cell type fraction for four Visium datasets (1 ≤ *k* ≤ 4) by-TD based unsupervised FE. Top left: *k* = 1, top right: *k* = 2, bottom left: *k* = 3, bottom right *k* = 4.

**Figure 22.**
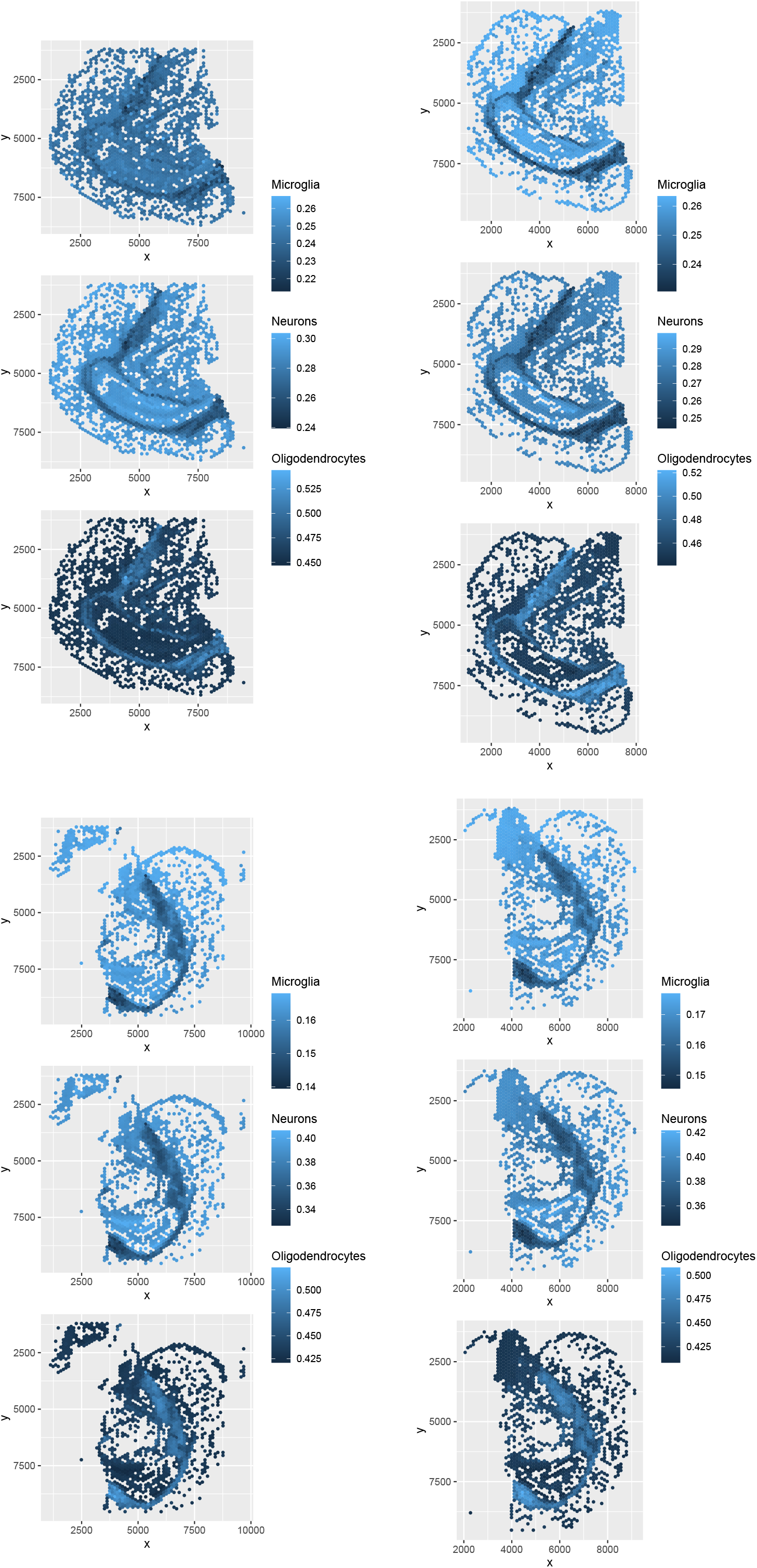
Spatial abundance of cell types for four Visium datasets (1 ≤ *k* ≤ 4) by-TD based unsupervised FE. Top left: *k* = 1, top right: *k* = 2, bottom left: *k* = 3, bottom right *k* = 4.

## 4. Discussion

Although one may wonder why we did not employ a more established benchmark in order to test the proposed method, we have already found that TD cannot compete with other SOTA when the established benchmark [18] is employed. Thus, we conclude that TD is superior to SOTA only when multiple minor cell types are present in the reference single-cell profiles. Although one might also wonder why we did not check all of the methods cited in the introduction for comparison, Because cell2location and RCTD are the top-ranked methods [18], checking the other inferior methods is time-consuming.

The reason why TD based unsupervised FE can reproduce spatial abundance (Fig. 22) well is as follows. First, before performing cell deconvolution, we confirmed that *v*_2*j*_ was associated with a reasonable spatial abundance (Fig. 2) and used the corresponding *u*_2*i*_ to select 277 genes. Then, only the expression of these 277 genes in the reference (single cell), 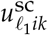, was used to predict the cell type fractions in individual spots. Such a procedure is possible only for TD based unsupervised FE and not for the other four methods. In addition, as shown in Table 2, spots that were unlikely to be correctly predicted were excluded (Fig. 22). This is another advantage of performing more accurate predictions. These two points allow TD based unsupervised FE to perform more accurate predictions.

## 5. Conclusions

In this paper, we demonstrated how TD-based unsupervised FE successfully performs deconvolution even when the reference single cell profile includes multiple minor cell types, for which four conventional methods fail. We also demonstrate that the identification of singular value vectors consistent with spatial cell type distribution may be the reason of the successful deconvolution.

## Author Contributions

Y. H. T. planned the study and performed the analyses. Y.-H.T. and T.T. evaluated the results and wrote and reviewed the manuscript. All authors have read and agreed to the published version of the manuscript. The conceptualization, data curation, analysis were performed by Y. H. T.

## Funding

This study was supported by a KAKENHI Grant (Number 24K15168).

## Institutional Review Board Statement

Not applicable

## Informed Consent Statement

Not applicable

## Data Availability Statement

All data analyzed in this study are from GEO and are publicly available.

### Code availability

https://github.com/tagtag/TDbasedUFE_deconvolution

## Acknowledgments

Not applicable

## Conflicts of Interest

We declare that the authors have no conflicts of interest.

## Disclaimer/Publisher’s Note

The statements, opinions and data contained in all publications are solely those of the individual author(s) and contributor(s) and not of MDPI and/or the editor(s). MDPI and/or the editor(s) disclaim responsibility for any injury to people or property resulting from any ideas, methods, instructions or products referred to in the content.

## Notes

### Competing Interest Statement

The authors have declared no competing interest.

### Summary of Updates

missing author family name missing author family name missing author family name missing author family name missing author family name missing author family name

